# Multiscale Segmentation using Hierarchical Phase-contrast Tomography and Deep Learning

**DOI:** 10.1101/2025.05.15.654263

**Authors:** Yang Zhou, Shahab Aslani, Yousef Javanmardi, Joseph Brunet, David Stansby, Saskia Carroll, Alexandre Bellier, Maximilian Ackermann, Paul Tafforeau, Peter D. Lee, Claire L. Walsh

## Abstract

Biomedical systems span multiple spatial scales, encompassing tiny functional units to entire organs. Interpreting these systems through image segmentation requires the effective propagation and integration of information across different scales. However, most existing segmentation methods are optimised for single-scale imaging modalities, limiting their ability to capture and analyse small functional units throughout complete human organs. To facilitate multiscale biomedical image segmentation, we utilised Hierarchical Phase-Contrast Tomography (HiP-CT), an advanced imaging modality that can generate 3D multiscale datasets from high-resolution volumes of interest (VOIs) at ca. 1 *µm*/voxel to whole-organ scans at ca. 20 *µm*/voxel. Building on these hierarchical multiscale datasets, we developed a deep learning-based segmentation pipeline that is initially trained on manually annotated high-resolution HiP-CT data and then extended to lower-resolution whole-organ scans using pseudo-labels generated from high-resolution predictions and multiscale image registration. As a case study, we focused on glomeruli in human kidneys, benchmarking four 3D deep learning models for biomedical image segmentation on a manually annotated high-resolution dataset extracted from VOIs, at 2.58 to ca. 5 *µm*/voxel, of four human kidneys. Among them, nnUNet demonstrated the best performance, achieving an average test Dice score of 0.906, and was subsequently used as the baseline model for multiscale segmentation in the pipeline. Applying this pipeline to two low-resolution full-organ data at ca. 25 *µm*/voxel, the model identified 1,019,890 and 231,179 glomeruli in a 62-year-old donor without kidney diseases and a 94-year-old hypertensive donor, enabling comprehensive morphological analyses, including cortical spatial statistics and glomerular distributions, which aligned well with previous anatomical studies. Our results highlight the effectiveness of the proposed pipeline for segmenting small functional units in multiscale bioimaging datasets and suggest its broader applicability to other organ systems.

## Introduction

Biological systems are organised across multiple 3D scales, from the macro scale of complete organs to the micro scale, including capillaries and functional units such as alveoli and nephrons. Functional units are considered the smallest unit of organs that perform a unique physiological function, and thus, studies on them provide a deeper fundamental understanding of how organs function Bidanta et al. [2025]. Segmentation of micro-scale functional units across entire organs enables analysis of functional relations and distributions relative to the whole organ, as well as variations across healthy human and pathological contexts.

However, such research is limited by the lack of 3D image datasets which simultaneously provide both high-resolution for micro-scale segmentation and whole organ coverage Pereira et al. [2016]. Although efficient multi-modal pipelines can provide such data, they are logistically complex, and the computational demand for accurate registration remains a challenging issue Paverd et al. [2024]. For example, histological sectioning can capture the 2D micro-scale functional units but does not image 3D structures Tyson et al. [2021]. Clinical imaging techniques, such as computed tomography (CT), magnetic resonance imaging (MRI), and ultrasound, effectively visualise human organs in 3D but are limited to macro-scale structures due to their resolution constraints Florkow et al. [2022].

Recently, these barriers have been overcome by the development of Hierarchical Phase-Contrast Tomography (HiP-CT) Walsh et al. [2021], a synchrotron-based X-ray technique, that can image intact human organs at ca. 20 *µm*/voxel down to micro-scale volume-of-interests (VOIs) at ca. 1 *µm*/voxel. Consecutive 3D scans from complete organs to high-resolution VOIs allow visualising and aligning tiny functional units across different scales.

Current studies on HiP-CT images, such as Jain et al. [2024], Yagis et al. [2024], Brunet et al. [2024], succeeded in 3D vasculature segmentation across multiple scales using manually annotated datasets. However, manually annotating HiP-CT data at different scales is time-consuming and labour-intensive. In the case of small functional units, it may not be possible to unambiguously generate manual annotations for small structures at lower resolutions (e.g., 20 *µm*/voxel). Moreover, scanning the complete organ at the highest possible resolution (e.g., 1 *µm*/voxel) is not routinely desirable and efficient due to the limitations of longer scanning, high X-ray radiation dose, and large data volumes. As a compromise, HiP-CT instead typically provides high resolution in smaller VOIs and lower-resolution overviews of whole organs. This multiscale structure makes HiP-CT an ideal modality for developing an automated multiscale pipeline to propagate segmentations from densely annotated high-resolution images to low-resolution whole organ images.

Addressing the limitations of manual annotation and resolution variability in HiP-CT demands robust segmentation techniques. Early biomedical image segmentation methods based on edge detection and region growing Azad et al. [2024] demonstrated good segmentation performance but were inefficient and struggled with blurry boundaries. Deep neural networks have provided significant advancements at independent single scales, varying from macro-scales organs (∼5*mm*/voxel) Gibson et al. [2018] to micro-scales functional units (under ∼ 1*mm*/voxel) Jiang et al. [2021]. Although several studies have explored segmentation on multiscale images, these typically generated the region-of-interests (ROIs) through resampling techniques such as pyramid pooling Al-Masni and Kim [2021] or scaling Srivastava et al. [2021]. Because these ROIs are not independently acquired but derived from the same dataset, they often suffer from pixelation artifacts and lack true multiscale fidelity.

To surmount these problems, we propose a multiscale biomedical image segmentation pipeline that leverages the resolution hierarchy of HiP-CT and deep learning to enable the analysis of micro-scale functional units across macroscale complete organs. The multiscale segmentation pipeline starts with training models on manually annotated high-resolution data and hierarchically propagates the segmentation capability to lower-resolution datasets with larger fields of view through multiscale correlative registration and fine-tuning with pseudo-labels. As a case study in this work, we focus on segmenting the glomerulus (plural: glomeruli), an approximately spherical-shaped capillary network of ca. 200 *µm* in diameter, which is situated within the cortex of the human kidney and plays a key role in blood filtration and autoregulation Singh Samant et al. [2023]. A single healthy kidney contains an estimated 330,000 to 1,400,000 glomeruli Basgen et al. [1994]. Prior glomeruli segmentation has primarily relied on 2D high-resolution (e.g., 0.85 *µm*/pixel) histological images, which limit 3D morphological analysis due to distortions and incomplete volumetric information Kannan et al. [2019]. To enable comprehensive 3D analysis of glomeruli, such as spatial distributions, on the complete human kidney imaged by HiP-CT, we manually annotated a high-resolution HiP-CT dataset (ca. 2.5 - 5 *µm*/voxel) of glomeruli and applied our multiscale segmentation pipeline to two different kidneys at ca. 25 *µm*/voxel.

In summary, the key contributions of this study are: (1) We present a 3D HiP-CT multiscale glomeruli segmentation dataset, including manual annotation on high-resolution VOIs from four intact human kidneys; (2) We propose a multiscale segmentation pipeline that can propagate the segmentation capability from annotated high-resolution HiP-CT VOIs to low-resolution intact organs volumes through multiscale registration and fine-tuning with pseudo-labels; (3) We benchmark four state-of-the-art 3D deep learning models: VNet Milletari et al. [2016], UNETR Hatamizadeh et al. [2022a], SwinUNETR Hatamizadeh et al. [2022b] and nnUNet Isensee et al. [2021] on high-resolution annotated data. The best-performing model is selected for correlative segmentation across scales; (4) We apply the full pipeline to two complete human kidneys from a 62-year-old donor without kidney diseases and a 94-year-old hypertensive donor, enabling downstream 3D morphological analysis of segmented glomeruli. The results are aligned with current anatomical studies.

All code is publicly available at https://github.com/UCL-MSM-Bio/2025-zhou-hipct-hierarchical-segmentation. git, and high-resolution annotated data can be accessed at https://doi.org/10.5281/zenodo.15397768.

## Related works

Biomedical image segmentation is important to generate pixel-wise (2D) and voxel-wise (3D) labels of distinct structures, potentially aiding clinical applications such as automated diagnosis Gao et al. [2021], Soomro et al. [2022]. Previous manual and semi-automated segmentation techniques are time-consuming and labour-intensive. With the advancement of deep neural networks, early automated methods aimed to solve 2D segmentation problems. Fully connected networks (FCN) Long et al. [2015] were the first end-to-end network that could generate direct and dense segmented predictions at an arbitrary input size. However, better FCN performance is limited by the depth of the network due to the gradient vanishing problem Hesamian et al. [2019]. To alleviate this, short skip connections Drozdzal et al. [2016], similar to residual connections, were introduced. FCNs also have difficulty preserving fine-grained details during upsampling without effective network structures. Therefore, Ronneberger et al. [2015] proposed a symmetric encoder-decoder structure, the U-Net, to propagate the context information in the upsampling stage. UNet++ Zhou et al. [2018] replaced the U-Net direct skip connections with nested and dense skip connections to enrich the feature maps from the encoder network to improve the segmentation performance.

Considering biomedical images are typically volumetric, and training a model on 2D slices only does not allow for the learning of 3D morphological features, V-Net Milletari et al. [2016] refined the 2D U-Net with 3D convolutional kernels and added residual connections in downsampling to accelerate convergence on 3D data. Since biomedical segmentation benefits from wide contextual information, non-local U-Net Wang et al. [2020] used an aggregation block based on a non-local attention mechanism Wang et al. [2018] to involve the global information without a deep encoder network. However, it is difficult to transfer a universal network structure across 3D biomedical images with different properties, like voxel spacing, and different modalities. Those properties illustrate the physical distances (spacing) and statistical differences (modalities) for the input images Zhou et al. [2019]. Thus, Isensee et al. [2021] proposed nnU-Net, an automated biomedical image segmentation pipeline, involving pre-processing, network structure configurations, and post-processing for any incoming dataset. To improve small object segmentation, omitted by U-shape network structures, KiU-Net Valanarasu et al. [2020] transferred the input to a higher spatial dimension and downsampled it to obtain the same-size output. Despite their success, these networks fail to capture long-range spatial dependencies and global context in challenging datasets.

Inspired by language transformers that can preserve long-range information, Vision Transformers (ViT) Dosovitskiy et al. [2020] allow training a Transformer network with image patches as tokens, achieving promising performances on classification tasks. TransUNet Chen et al. [2021] firstly incorporated the Transformer self-attention block into CNN as a hybrid CNN-Transformer network, showing the potential on the 2D biomedical image segmentation task. For 3D image segmentation, UNETR, developed by Hatamizadeh et al. [2022a], directly applied the Transformer as an encoder and used 3D patches as inputs. SwinUNETR Tang et al. [2022] integrates the hierarchical encoder from Swin Transformer to achieve better segmentation performances.

Although the above networks are promising for HiP-CT segmentation, pseudo-label propagation between different resolutions requires fine-tuning techniques. A previous study in Mahbod et al. [2020] showed that re-training the network on different scales transferred the classification performance of skin lesions across different resolutions. MorphHR Wei et al. [2022] fine-tuned the network with morphed layers and successfully applied the model that is trained on natural images to high-resolution mammogram images for classification tasks. However, fine-tuning a network for segmentation tasks and multiscale biomedical images with different voxel sizes remains a research gap.

## Data

HiP-CT generates multiscale 3D images of intact human organs from ca. 20 *µm*/voxel to zoomed VOIs at ca. 1 *µm*/voxel. This hierarchical feature enables morphological studies on small functional units, e.g., glomeruli in the diameter size range of 100 *µm* to 200 *µm*, and statistical analysis across whole organs. As HiP-CT creates nested VOIs of increasing resolution without the need to section the organ physically, alignment between different resolutions is relatively straightforward. Fig. 1 (A) shows registered slices in different resolutions of one kidney, and Fig. 1 (B) depicts the 3D visualisations of registered VOIs in this kidney.

**Figure 1.**
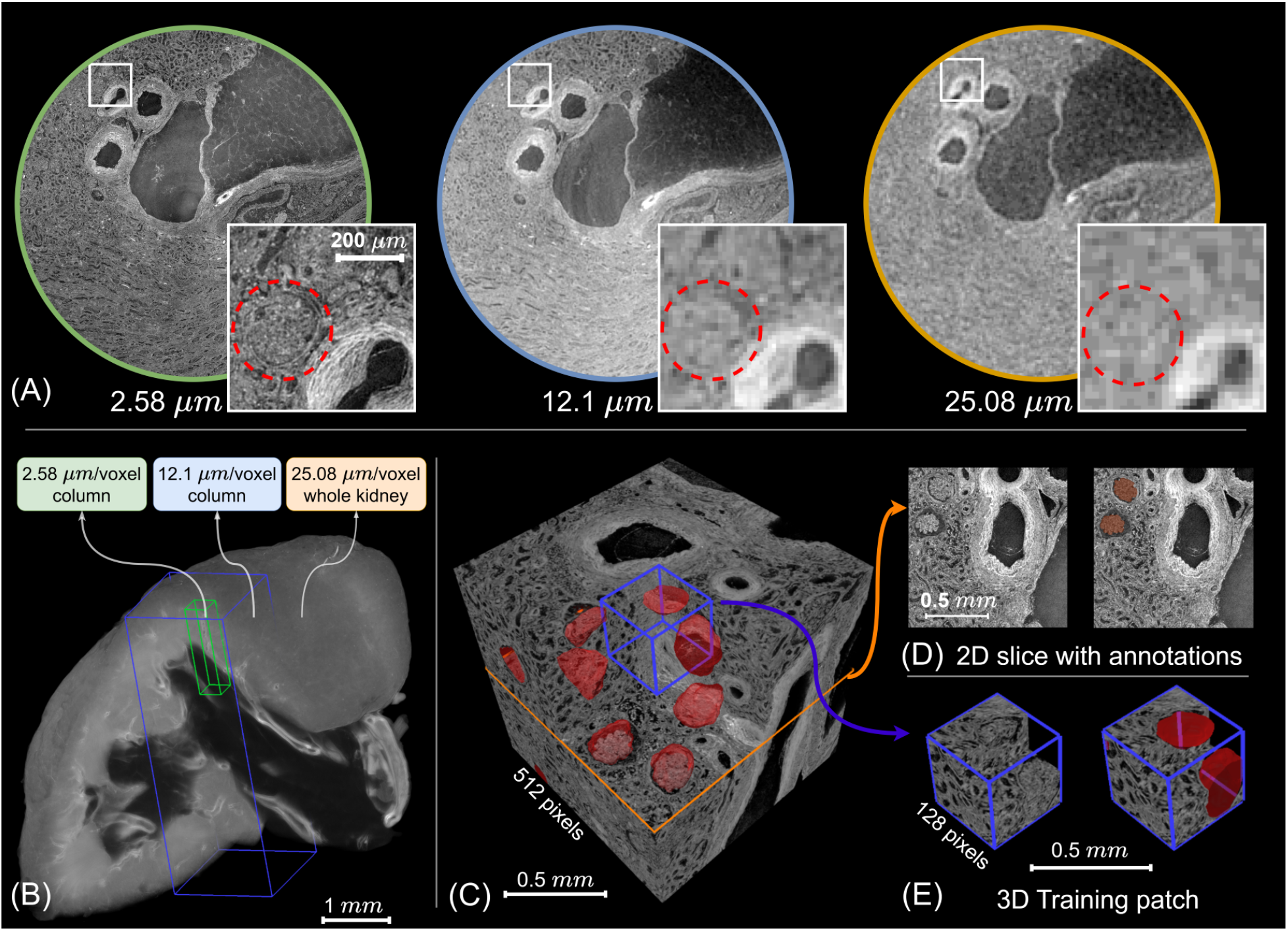
Multiscale HiP-CT dataset for glomeruli segmentation from the sample of LADAF-2020-17 left kidney. (A) Visualisation of 2D slices from the same region at three different resolutions: 2.58 *µm*/voxel Tafforeau et al. [2021a], 12.1 *µm*/voxel Tafforeau et al. [2021b], and 25.08 *µm*/voxel Tafforeau et al. [2021c]. Inset squares highlight the same glomerular region across resolutions. (B) 3D rendering of the full kidney at 25.08 *µm*/voxel, with cuboids indicating the locations of two selected higher-resolution volumes at 12.1 *µm*/voxel and 2.58 *µm*/voxel. (C) Manually annotated training volumes in dimension of 512^3^, where glomeruli are marked in red. (D) Example 2D slice of the training cube with glomeruli annotated in red. (E) Cropped 3D Training patches in dimension of 128^3^, used as inputs for neural network training.

The multiscale HiP-CT glomeruli dataset involves three different resolutions: high resolutions of ca. 2.58 to 5.2 *µm*/voxel, intermediate resolution of ca. 12 *µm*/voxel, and low resolution of ca. 25 *µm*/voxel. To reduce the data size and computational resource needs, the high-resolution and intermediate-resolution data were binned by a ratio of two. Four different kidneys from three donors were used, as shown in Table 1, prepared in ethanol before being scanned at the European Synchrotron Radiation Facility (ESRF) with HiP-CT (see Methods for details). Manual annotation was conducted on 40 cubes with dimensions of 512^3^ from high-resolution volume-of-interests (VOIs) of those four kidneys, varying from 2.58 *µm*/voxel to 5.2 *µm*/voxel (binned by 2 to reduce the data size). To annotate the glomeruli on the high-resolution VOIs, the annotators evaluated several adjacent slices to label them according to obvious 3D morphological features.

**Table 1.** Information and statistics of HiP-CT glomeruli segmentation dataset for training based on 9:1 train-test split across 4 samples.

| Samples | Information | Training |  | Testing |  |
| --- | --- | --- | --- | --- | --- |
|  |  | Cubes | Patches | Cubes | Patches |
| 5 $\mu\text{m}$ S-20-28 | Male, aged 84, prepared with ethanol | 18 | 831 | 2 | 119 |
| 5.2 $\mu\text{m}$ LADAF-2021-17 Left Kidney C. L. Walsh et al. [2024a] | Male, aged 63, prepared with ethanol | 6 | 187 | 1 | 10 |
| 5.2 $\mu\text{m}$ LADAF-2021-17 Right Kidney C. L. Walsh et al. [2024b] | Male, aged 63, prepared with ethanol | 8 | 226 | 1 | 13 |
| 2.58 $\mu\text{m}$ LADAF-2020-27 Left Kidney Tafforeau et al. [2021a] | Female, aged 94, prepared with ethanol | 3 | 86 | 1 | 26 |
| Total | - | 35 | 1330 | 5 | 168 |

Dataset statistics are shown in Table 1. Due to computational resource limitations, the manually annotated cubes of 512^3^ were cropped into non-overlapping 3D patches of 128^3^ voxels for training. Fig. 1 (C) shows one of the labelled cubes in both 2D slices and 3D patches. We aimed to follow a ratio of 9:1 to split the train-test dataset based on the original cubes of dimensions 512^3^. Considering the imbalance in the number of cubes annotated for each kidney, at least one cube was selected for the test dataset from every kidney. The statistics for data splits are shown in Table 1.

## Methods

### Data aquisition

HiP-CT is an X-ray phase-contrast propagation technique using the Extremely Brilliant Source (EBS) from the European Synchrotron Radiation Facility (ESRF). Prior work Walsh et al. [2021], Brunet et al. [2023] describes the sample preparation and imaging procedure.

Four human kidneys were obtained with ethical approvals from three donors, LADAF-2020-27, LADAF-2021-17, and S-20-28. LADAF-2021-17 left kidney, LADAF-2021-17 right kidney, and LADAF-2020-27 left kidney were collected from the donors who had consented to body donation to the Laboratoire d’Anatomie des Alpes Françaises (LADAF) before death. The kidney of S-20-28 was obtained after a clinical autopsy at the Hannover Institute of Pathology at Medizinische Hochschule, Hannover (Ethics vote no. 9621 BO K 2021). The transportation and imaging protocols received approval from the French Health Ministry. HiP-CT scan parameters and relevant tomographic information are presented in the Supporting Information S1 Text, section 1.

### Multi-resolution segmentation pipeline

The proposed multiscale segmentation pipeline took annotated HiP-CT data (Fig. 2.A) as inputs and generated predictions of glomeruli labels for the complete VOI (Fig. 2.E) at the same resolution as the inputs, iteratively propagating the segmentation capability from high-resolution VOIs to low-resolution complete organs. Each hierarchical cycle included five steps: data pre-processing (Fig. 2.B), training/fine-tuning the deep neural network (Fig. 2.C), prediction post-processing (Fig. 2.D), multiscale registration (Fig. 2.F), and pseudo-labelling lower-resolution data (Fig. 2.G), which are described in the following sections.

**Figure 2.**
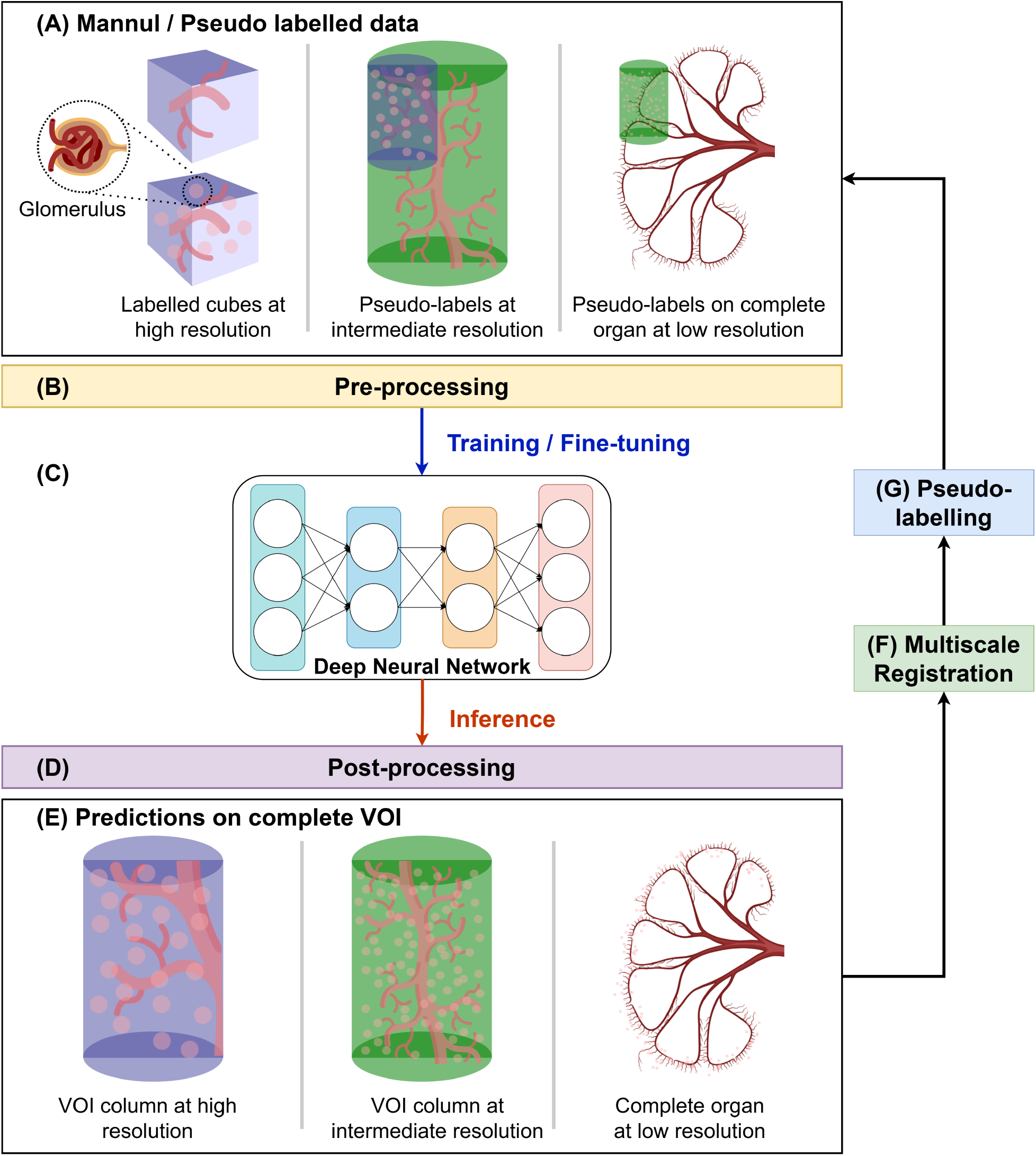
Overview of the proposed multiscale segmentation pipeline using glomeruli as a case study. This pipeline enables segmentation of tiny functional units across multiple resolutions of HiP-CT data. (A) Input data at multiple HiP-CT resolutions, with corresponding manual annotations or pseudo-labels; (B) Pre-processing steps, including contrast enhancement and patch extraction, to improve image quality and address memory constraints; (C) Training or fine-tuning of deep neural networks using the labelled HiP-CT data; (D) Post-processing to eliminate false positives based on intensity and morphological criteria; (E) Final post-processed predictions on the complete HiP-CT volume of interest (VOI); (F) Multiscale registration aligning higher-resolution VOIs with lower-resolution scans; (G) Generation of pseudo-labels for low-resolution data by propagating high-resolution predictions through the registration transforms.

To segment the glomeruli on high-resolution data and generate the pseudo-labels on lower-resolution data, the pipeline starts to train the deep neural network with manually labelled cubes extracted from high-resolution volumes. Since the high-resolution volume covers only a small region in the intact kidney, directly downsampling the predictions to the intact human organ at the lowest resolution could not provide enough pseudo-labels to fine-tune the models. Introducing an intermediate resolution is essential to increase the amount of pseudo-labels. Therefore, in this work, the multi-resolution glomeruli segmentation pipeline involves three stages across three resolutions.

### Data pre-processing

As the voxel intensity of HiP-CT data is non-quantitative and can vary substantially between samples due to imaging setup, anatomical variation, sample preparation, etc., it is essential to normalise datasets across different samples. To do this, we applied contrast-limited adaptive histogram equalisation (CLAHE) Pizer et al. [1987] to each sample, which decreases the noise amplified in the near-contrast regions (see S1 Text, section 3).

The original HiP-CT images form large datasets (typically 250 GB - 1 TB per volume). Training the deep neural networks on such big datasets is challenging, and following preliminary experimentation, we found that conversion of the data from 16-bit to 8-bit after application of CLAHE reduced the memory requirements and allowed for better optimisation of the pipeline, leading to better overall segmentation outputs. After that, division of the original labelled cubes (512^3^ voxels) into non-overlapping cubes of 128^3^ was performed. This size enabled training a model with a larger batch size while being able to keep a whole glomerulus in the volume of interest. To keep the consistency and the network structure in the fine-tuning, the same preprocessing techniques were applied to the lower-resolution data in the pipeline.

### Training with manual labels

Since HiP-CT generates 3D isotropic data, we tested four 3D segmentation deep learning models: VNet, UNETR, SwinUNETR, and nnU-Net for the first stage of high-resolution data with manual labels.

The performance of these four models was evaluated by the average Dice score, a metric measuring the overlap area between predictions and ground truths, using 5-fold cross-validation. The highest performing network was selected as the baseline model to then perform the correlative segmentation at lower resolutions. The models were developed with PyTorch Ansel et al. [2024] 2.0.1, and all the experiments were performed on four Tesla V100 GPUs, and the inference was on one NVIDIA RTX A6000 GPU.

### Post-processing

In this pipeline, post-processing was crucial for eliminating false positives before registration and propagating pseudolabels to lower-resolution data. It improved the quality of the automated labels used in fine-tuning the network.

After training, we applied inference of the model to the complete VOI from which the training cubes had been extracted. However, the predictions showed a large number of false positives. To remove these false positives, we generated instance segmentations where each predicted glomerulus has associated intensity and morphological properties. We then selected several representative parameters, such as intensity variance, roundness, and size, and generated histograms of these properties to filter the outliers.

Considering the organ shrinkage due to dehydration during preparation, the minimum radius of a glomerulus is set to 62 *µm* (volume= 1 × 10^6^*µm*^3^), which is close to the smallest measurement from MRI in Beeman et al. [2014] of 72 *µm* (volume= 1.6 × 10^6^*µm*^3^). Apart from this fixed minimum size, other thresholds for the selected parameters are automatically set based on the percentiles of the aggregate data for all glomeruli. From these threshold bounds, Latin Hypercube Sampling (LHS) McKay et al. [2000] was then used to explore the parameter space to optimise the combination of post-processing. 20 parameter sets were generated through LHS, and each parameter set was evaluated using Dice scores on all labelled cubes and an additional empty cube containing only fat, where no glomeruli should be present, and where the majority of the false positives were found. We also propose an additional evaluation metric, an instance Dice score *Dice*_*I*_ as shown in Eq. 1, to evaluate the effectiveness of thresholding.

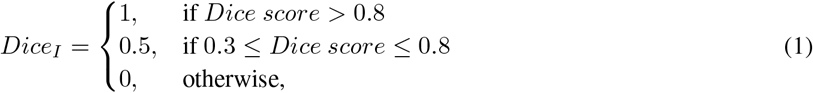

where the *Dice* is the normal Dice score. Since precise predictions of the blurry boundaries of glomeruli in low-resolution data are difficult, we propose the *Dice*_*I*_ to indicate a higher overlap rate between predictions and ground truths, which helps perform density and distribution studies of the glomeruli on the low-resolution data.

The properties and the threshold criteria are listed in Table 2 in the Results Section.

**Table 2.** Post-processing parameters and corresponding Dice scores (before and after post-processing) for predictions on LADAF-2020-27 left kidney scans at different resolutions.

| Parameters | 2.58 $\mu\text{m}/\text{voxel}$ prediction | 12.1 $\mu\text{m}/\text{voxel}$ prediction | 25.08 $\mu\text{m}/\text{voxel}$ prediction |
| --- | --- | --- | --- |
| Variance lower bound | 114770.713 | 179663.370 | N/A |
| Variance upper bound | 3069051.733 | N/A | N/A |
| Roundness | 0.709 | 0.682 | N/A |
| Size (radius in $\mu\text{m}$) | 62 | 62 | 62 |
| Density | N/A | $W^{[1]}=128, D^{[2]}=3$ | $W=64, D=3$ |
| Mask | N/A | N/A | Cortex mask applied |
| Dice scores (before/after) | 0.734 / 0.934 | 0.475 / 0.974 | 0.786 / 0.788 |
| Instance Dice scores (before/after) | 0.754 / 0.954 | 0.493 / 0.992 | 0.785 / 0.785 |

[1] windows size
[2] density of neighbourhood

### Multiscale registration

Intact human organs at lower resolution are scanned first, followed by the high-resolution VOIs. Therefore, the multi-resolution VOIs must be registered to the complete organ volume to align the features during correlative model fine-tuning using higher-resolution pseudo-labels.

The registration was implemented utilising SimpleITK Lowekamp et al. [2013] and Mattes Mutual Information Mattes et al. [2001, 2003] as a similarity metric over optimisation, between the lower-resolution fixed volume and the higher-resolution moving volume. The process started with two manually selected common points in each solution as a fixed centre. The common points are identical features among multi-resolution data recognised by experienced experts. With the fixed centre, the registration process started with finding the *z*-rotation angle between the high-resolution and the low-resolution volumes. The process involves two consecutive exhaustive searches - 360 degrees with 2 degrees as a step and 5 degrees with 0.1 degrees as a step. By searching on a large range and then a smaller range, the registration process can quickly optimise to find the z-rotation angles. After that, the *xy*-plane rotation angle and scaling factor are optimised by the Limited-memory Broyden–Fletcher–Goldfarb–Shanno (LBFGSB) algorithm with 2000 iterations. The registration results are presented in the S1 Text, Section 4.

### Pseudo-labelling and fine-tuning

After obtaining registration matrices, the post-processed predicted labels of the higher-resolution volume were registered to the lower-resolution data. Based on the locations of the predicted labels, the low-resolution VOI and their labels were cropped into cubes with size 128^3^ as a training dataset, consistent with the cube size used to train at higher resolution data. Then, the trained model was fine-tuned on the new data, adapting it to lower-resolution data. The hyperparameters of the model used for fine-tuning are the same as the hyperparameters used for the high-resolution training stage.

For the lowest-resolution data, ca. 25 *µm*/voxel, the glomeruli features are subtle in lower-resolution data due to the high voxel size. To obtain the best segmentation performance with the pseudo-labels, we explored different training strategies, such as training from scratch and changes in learning rates, which are presented in the Supporting Information S1 Text, Section 6.

### Deep neural network optimisation

The deep neural networks were optimised by an ensemble cost function *L* of a cross-entropy loss *L*_*CE*_ and a weighted Dice loss *L*_*Dice*_ for glomeruli segmentation as shown in Eq. 2.

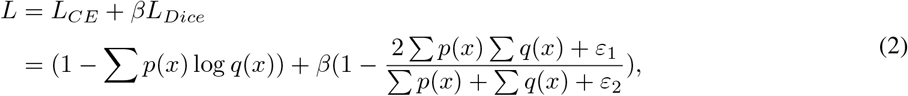

where *p*(*x*) and *q*(*x*) are the prediction and ground truth at a given voxel coordinate *x*, and the sums are over all voxels in a patch. *ε*_1_ and *ε*_2_ are small constant values, set to 1 × 10^−5^ as default, to avoid division by zero in the Dice loss. *β* is the weight of the Dice loss relative to the cross-entropy loss, and is set to 1 by default.

Cross-entropy loss is widely used in segmentation tasks to quickly train the network based on pixel-wise error and stable gradients. However, it tends to place larger weights on the larger objects while neglecting small ones Yeung et al. [2022]. Considering that the glomeruli segmentation consists of two classes (glomeruli or not-glomeruli) with the glomeruli class appearing sparsely, we added the additional Dice loss with a weight *β* to the overall loss. The Dice loss measures the overlap areas between predictions and labels, and is inherently designed for imbalanced classes. Therefore, our cost function enables the networks to learn voxel-wise and regional overlap information.

## Results

### Glomeruli segmentation on high-resolution labelled data

The performance of the four initial network architectures was tested on the highest resolution data and compared using Dice scores across 5-fold cross-validation through training, validation, and test datasets. Figure 3 (A) shows the average Dice scores and their standard deviations revealed by the error bar after training each model with 150 epochs. After that, the Dice scores of the test dataset were calculated from the best models of each method.

**Figure 3.**
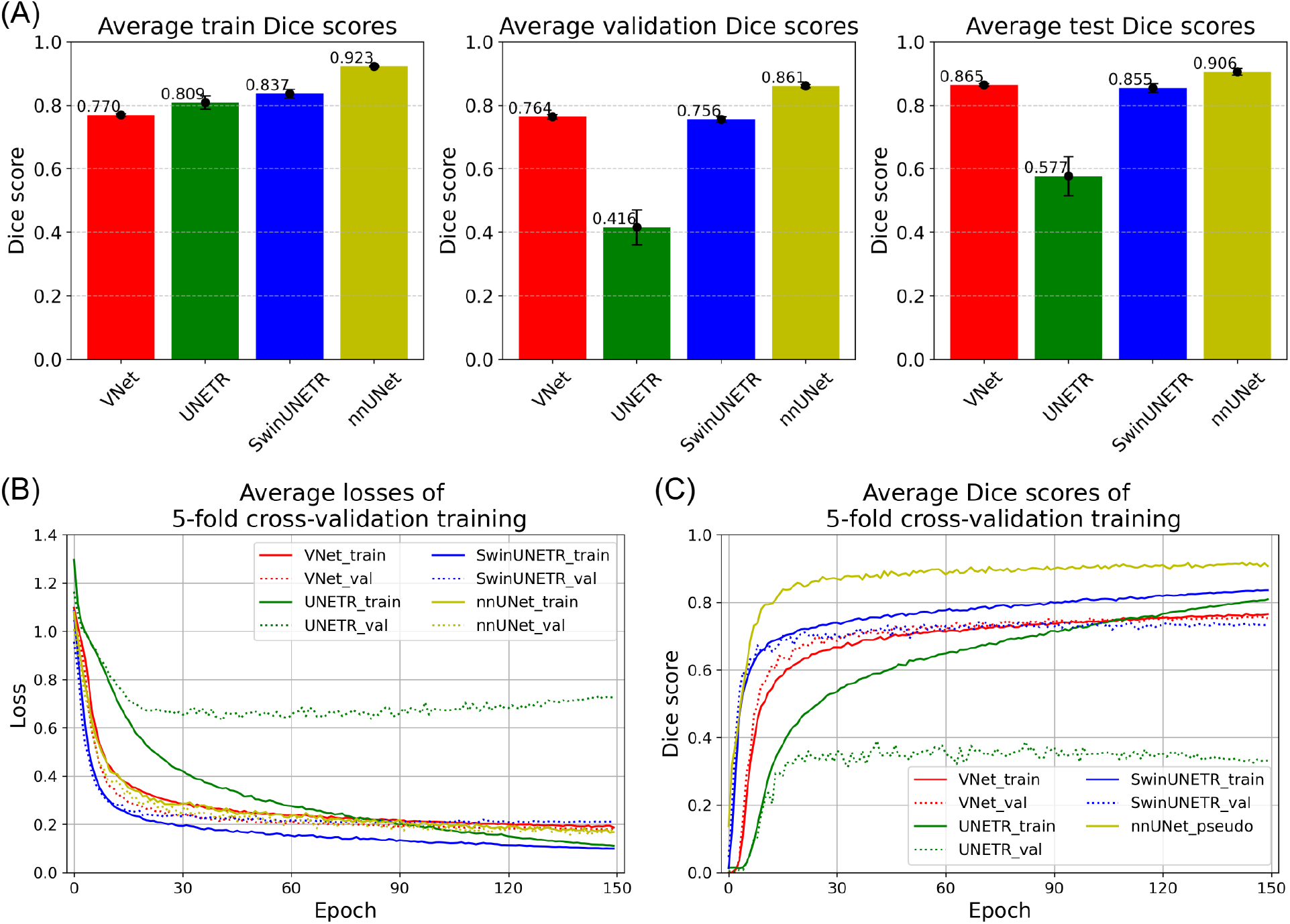
Performance of different models trained on high-resolution manually annotated data. Average training losses and Dice scores from four 3D segmentation models: VNet, UNETR, SwinUNETR, and nnUNet, evaluated using 5-fold cross-validation. (A) Average Dice scores on the training, validation, and test sets for each model, with error bars indicating inter-fold variance; (B) Training loss curves showing the average losses on the training and validation sets over epochs; (C) Average Dice scores for each model during training (nnUNet only reported the pseudo Dice score after a moving average applied).

As shown in Fig. 3, during training, nnUNet had the highest average Dice score of 0.923, followed by SwinUNETR of 0.837, UNETR of 0.809, and VNet of 0.770. However, VNet achieved a higher Dice score than the Transformer models in validation. As for the test dataset, we performed sliding windows inference using the best models of each method on the complete annotated data in the size of 512^3^, without cropping them to the size of training and validation patches of 128^3^. This improved the segmentation performance compared to the Dice scores of the validation data.

Fig. 3 (B) presents the average losses and (C) shows the average Dice scores on the training (solid lines) and validation (dotted lines) datasets of each model during 5-fold cross-validation training. As training progressed, both the training and validation losses for nnUNet and VNet decreased. However, for UNETR and SwinUNETR, the training losses decreased while the validation losses increased slightly at the end. Similar patterns were observed in the rise of the Dice scores. These indicated that the UNETR and SwinUNETR experienced the over-fitting problem, achieving better training performance on the training dataset but performing worse on the unseen data.

Fig. 4 visualises several validation samples from fold 0 with their ground truths and predictions from the four different methods. From these images, we saw that VNet, SwinUNETR and nnU-Net can predict smooth glomeruli boundaries similar to the annotations, but UNETR had difficulty learning the correct boundary features. This negatively impacted the UNETR performance and resulted in low Dice scores. nnU-Net outperformed other methods because fewer false positives were predicted, compared to other methods with plenty of false positive predictions, such as the first sample for UNETR, the second sample for SwinUNETR, and the fourth sample for VNet.

**Figure 4.**
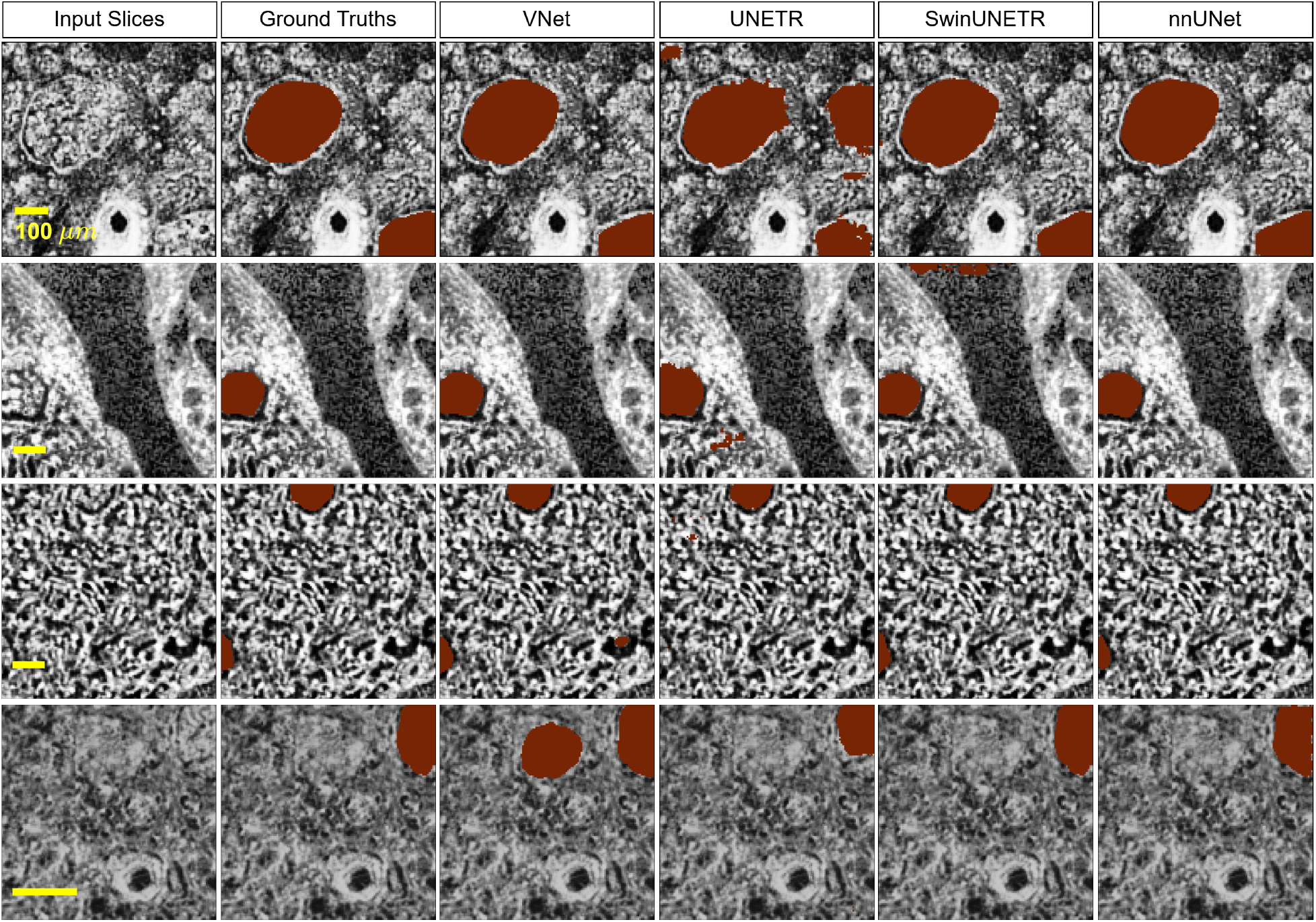
Ground truths from manual annotations and predictions (both in red) from each model in fold 0 on high-resolution data, shown in 2D slices. The 2D slices were sampled from validation cubes of each kidney; from top row down: 5 *µm* S-20-28, 5.2 *µm* LADAF-2021-17 left kidney, 5.2 *µm* LADAF-2021-17 right kidney, and 2.58 *µm* LADAF-2020-27 left kidney. The slices are of size 128^2^ and pre-processed by CLAHE.

### Post-processing effects

We also proposed an automated post-processing pipeline to remove the outliers in the predictions and obtain clean pseudo-labels for the lower-resolution data. The nnUNet with the best performance on the annotated high-resolution cubes was selected as a benchmark method for correlative glomeruli segmentation, but its predictions when performing inference on the whole VOI of HiP-CT data can be clearly seen to have a large number of false positives (see S1 Text section 5), primarily falling into three areas within the kidney: fat, blood clots, and tubular structures, as shown in Fig. 5.

**Figure 5.**
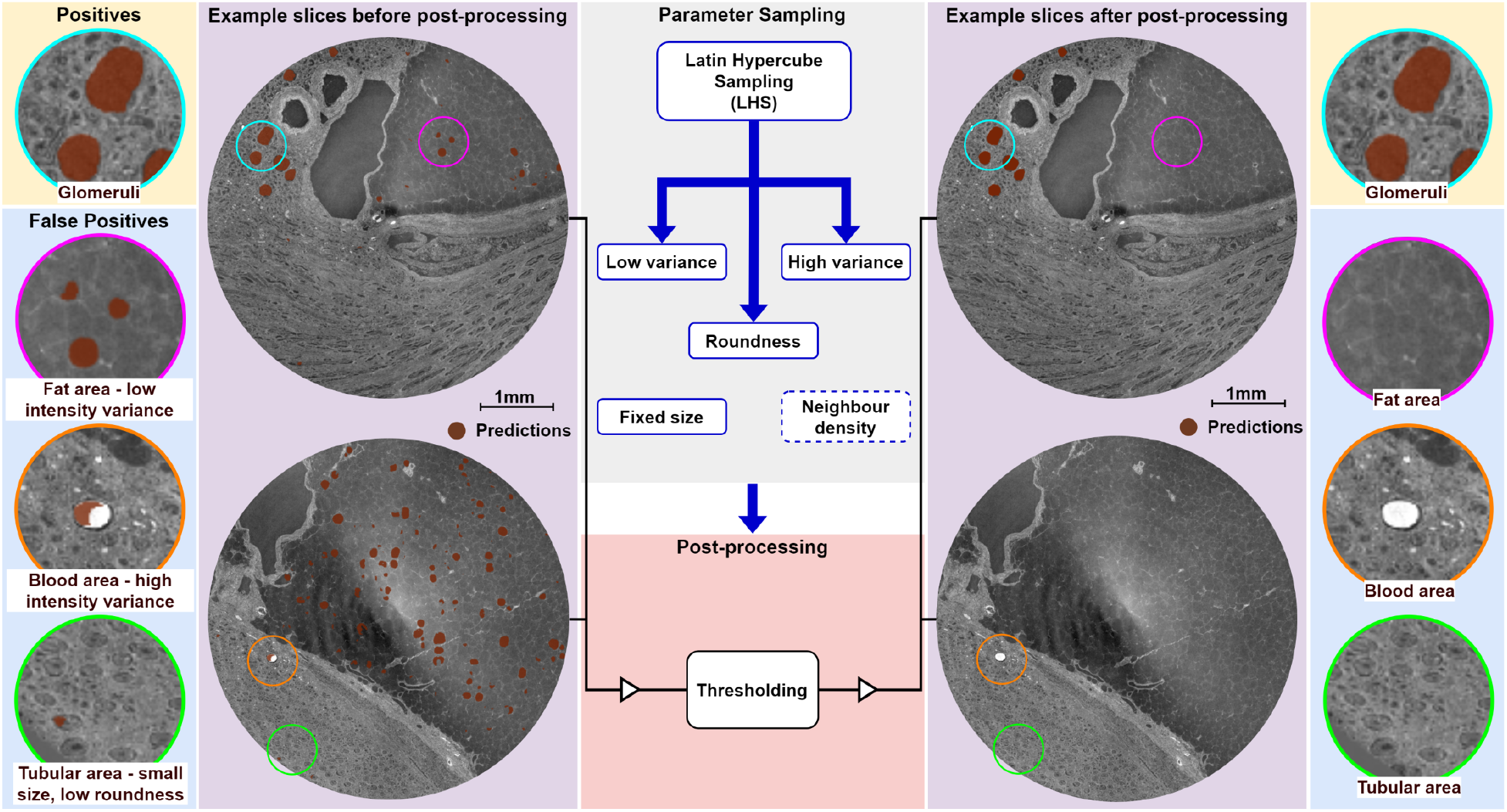
Post-processing to reduce false positives in glomeruli segmentation. False positives occurring in fat, blood clots, and tubular regions are removed using threshold-based filtering. Optimal thresholds were determined via Latin Hypercube Sampling (LHS). Representative examples are taken from the LADAF-2020-27 left kidney dataset.

During post-processing, parameter thresholds listed in Table 2 were sampled using LHS to eliminate outliers. To assess the performance of the sampled thresholds, we computed the Dice scores and instance Dice scores across all training cubes as well as in specific glomeruli-free cubes for both high and intermediate-resolution data. However, for the lowest-resolution data of complete organs, only the training cubes were utilised due to the larger VOIs with enough content (see S1 Text section 2).

A consistent minimum size threshold of 62 *µm* in radius was applied across all resolutions. For data with high contrast between the kidney cortex and other structures against the background, thresholds for intensity variance and roundness were employed (at high and mid resolutions). In contrast, for the data with lower contrast (mid and low resolution overviews), neighbourhood density was used due to the sparse distribution of false positives compared to true positives. Additionally, a cortex mask (manually annotated) was applied to whole-kidney predictions to eliminate false positives occurring in the medulla and hilum regions, where glomeruli are absent. After post-processing, Dice score and instance Dice scores were highly improved for high and intermediate-resolution data, while slightly improved for the low-resolution complete organ data.

### Glomeruli segmentation on lower-resolution data

After training on high-resolution data with manual annotations and post-processing, nnUNet performed best and was used for correlative lower-resolution data. This section reports the results of LADAF-2020-27 left kidney at 12.1 *µm*/voxel and 25.08 *µm*/voxel.

To fine-tune the models, the predictions were registered, and the corresponding VOI in the lower-resolution data was cropped as pseudo-labelled training data. The correlative fine-tuning followed 5-fold cross-validation and the same patch size of 128^3^. As the glomeruli were small and sparsely distributed in the image, the training data could include many cubes without glomeruli. Those empty cubes could bias the model training process. Therefore, at the intermediate resolution(12.1 *µm*/voxel) where the glomeruli can still be clearly identified, we only kept 1.05% of the empty cubes after cropping. However, for the data at 25.08 *µm*/voxel that has unclear glomeruli boundaries, we found that removal of the empty cubes was not enough to train a model with better performance. Accordingly, instead of keeping empty cubes, we kept 0.7% of the cubes with label volume densities lower than 1%. The average number of patches used in fine-tuning is shown in Table 3.

**Table 3.** Training samples and average Dice scores of correlative 5-fold training on lower-resolution data of LADAF-2020-27 left kidney with pseudo-labels.

| Resolutions | 12.1 $\mu\text{m}$ | | 25.8 $\mu\text{m}$ | |
| --- | --- | --- | --- | --- |
|  | Train | Val | Train | Val |
| Average number of patches | 204 | 51 | 1054 | 263 |
| Average of Dice scores | 0.950 | 0.949 | 0.775 | 0.796 |
| Standard deviation of Dice scores | 0.001 | 0.006 | 0.002 | 0.002 |

The model was fine-tuned with 1000 epochs on the 12.1 *µm*/voxel data and 1500 epochs on the 25.08 *µm* data, to reach the training plateau, as shown in Fig. 6. For fine-tuning on the 25.08 *µm*/voxel data, an ablation study for the learning rate was performed, and it was found that by continuing to train 500 more epochs after 1000 epochs with a starting learning rate of 0.00074, performance was improved (see S1 Text, Section 6, for details on the training explorations). As in Table 3, the model fine-tuned on the 12.1 *µm*/voxel data achieves an average Dice score of 0.950 on the training dataset, and 0.949 on the validation dataset. The model fine-tuned on the 25.08 *µm*/voxel data achieves scores of 0.775 and 0.796, respectively.

**Figure 6.**
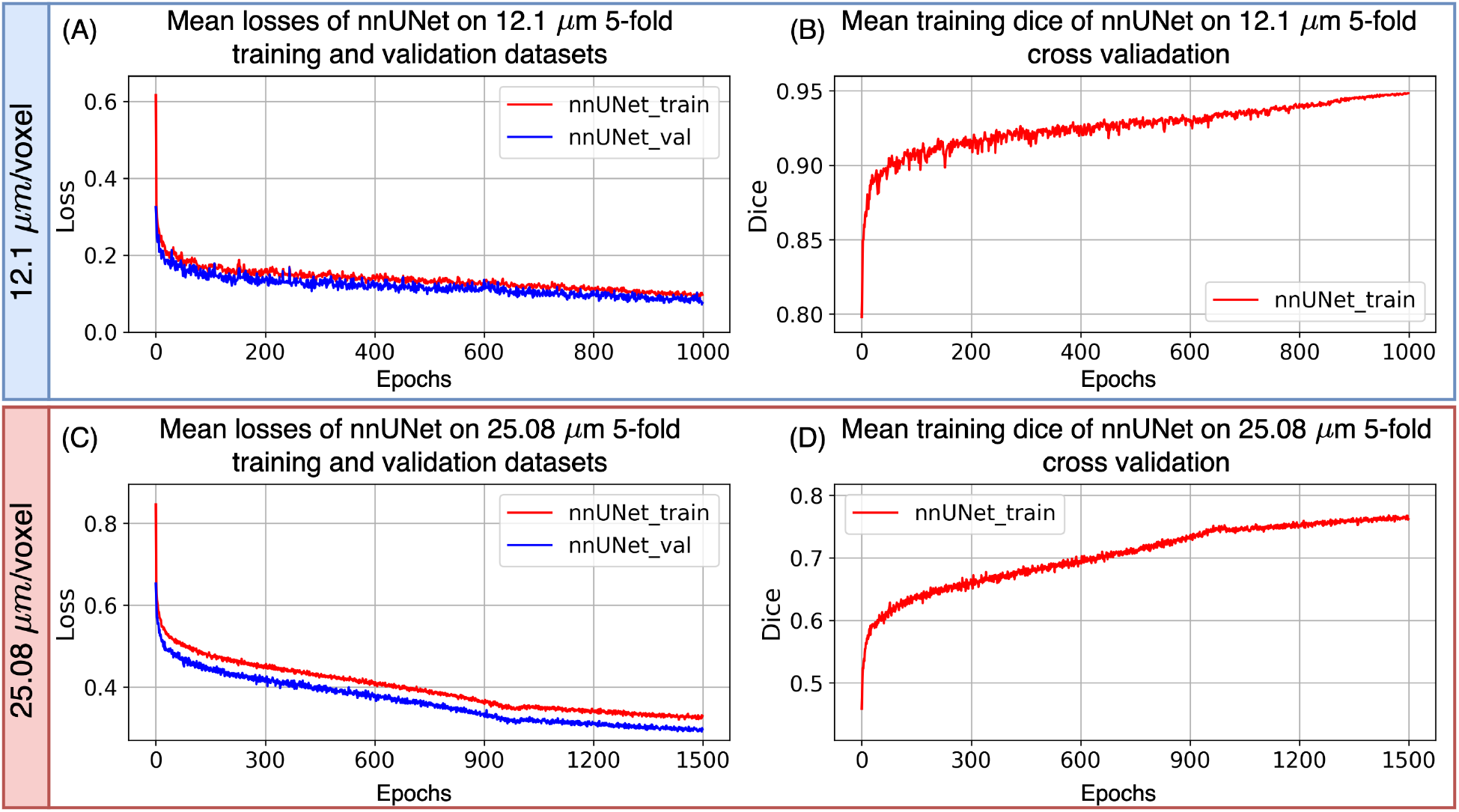
Correlative training on lower resolution data of LADAF-2020-27 left kidney with pseudo-labels. (A) and (B) show the mean losses and mean Dice scores from 5-fold cross-validation training at the intermediate resolution of 12.1 *µ*m/voxel. (C) and (D) present the same metrics for the lowest resolution data of 25.08 *µ*m/voxel.

### Morphological analysis

Our pipeline enables the morphological analysis of glomeruli across the whole human kidney. Having trained the model on high-resolution data, we applied the pipeline to two human kidneys: LADAF-2020-27 left kidney from a female donor aged 94, and LADAF-2021-17 right kidney Rahmani et al. [2024] from a male donor aged 64. The training details of LADAF-2021-17 right kidney are presented in the S1 Text Section 7. The female donor suffered from hypertensive heart disease Nemtsova et al. [2024], micro-crystalline arthritis (gout) Stamp et al. [2021], atrial fibrillation Kiuchi [2018], and right cerebellar stroke Zhao et al. [2020], all of which can directly impact glomerular functionality. No kidney-related diseases were observed in the male donor. These two kidneys could provide comparable analysis across different ages, sexes, and healthy statuses.

As shown in Table 4, the total and cortical volumes of the female donor’s kidney were approximately 70 *cm*^3^ and 40.6 *cm*^3^, respectively, while those of the male donor’s kidney were approximately 136 *cm*^3^ and 88.6 *cm*^3^, indicating a notable difference in organ size, potentially attributable to donor age, sex, and health status Dabers et al. [2024].

**Table 4.** Kidney details for morphological analysis.

| Kidneys | Sex | Age | Kidney volume ($\text{cm}^3$) | Cortex Volume ($\text{cm}^3$) | Number of glomeruli |
| --- | --- | --- | --- | --- | --- |
| LADAF-2020-27 Left | F | 94 | 69.74 | 40.56 | 231,179 |
| LADAF-2021-17 Right | M | 62 | 136.05 | 88.64 | 1,019,890 |

The predictions of glomeruli across the entire human kidneys, as shown in Fig. 7 (A.1 and B.1), facilitated anatomical evaluation, which is important as their location and size are linked to functional heterogeneity in filtration and susceptibility to disease across different renal regions Ikoma et al. [1990].

**Figure 7.**
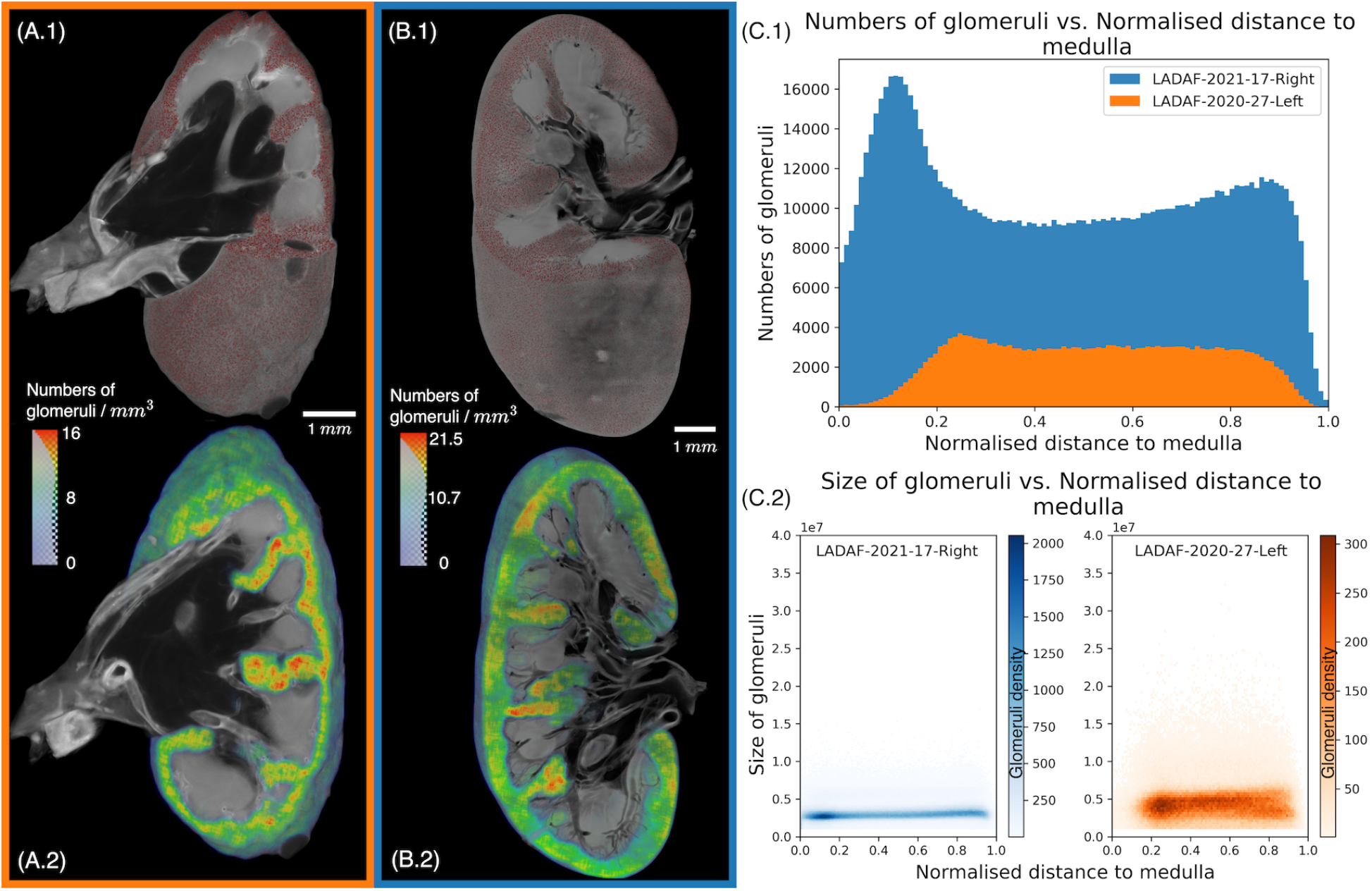
Glomeruli morphological analysis on the complete kidneys. (A.1) and (A.2) illustrate 3D rendering of predicted glomeruli and the density calculated by a mean kernel of the complete kidney (LADAF-2020-27-Left, Female, aged 94). Similarly, (B.1) and (B.2) are for LADAF-2021-17-Right, Male, aged 62. (C.1) depicts the number of glomeruli relative to the distance (normalised to [0,1]) to the medulla. (C.2) shows the glomeruli size relative to the distance (normalised to [0,1]) to the medulla.

Firstly, by convolving a mean kernel of size 31^3^ on the predictions, we generated the density maps as shown in Fig. 7 (A.2 and B.2) to display the glomeruli distributions within the kidney cortex. Additionally, the spatial distribution of glomeruli within the cortex is presented in Fig. 7 (C.1). The distances of glomeruli to the medulla and kidney capsule were calculated by the 3D Euclidean distance. After that, the distances to medulla were normalised to the range of [0, 1], as in Eq. 3 for each kidney, to better compare two kidneys of different size.

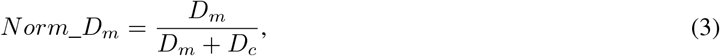

where *Norm*_*D*_*m*_, *D*_*m*_, and *D*_*c*_ denote the normalised distance to medulla, distance to medulla and distance to the capsule, respectively. For both kidneys, the number of glomeruli as a function of their relative distance to the medulla reveals a bimodal distribution, with prominent peaks near both the corticomedullary boundary and the outer capsule. This pattern suggests regional clustering of glomeruli that may reflect developmental or physiological zonation within the cortex Schaub et al. [2023].

Secondly, the 2D histograms in Fig. 7 (C.2) illustrate the distribution of glomerular volumes as a function of their relative distance to the medulla. The mean glomerular volume was 3.13 × 10^6^ ± 1.08 × 10^6^ *µm*^3^ in the healthy male kidney and 4.52 ×10^6^ ± 2.16 ×10^6^ *µm*^3^ in the female hypertensive kidney. Despite the overall uniformity in volume distribution across the cortex, segmenting the cortex into three zones—inner (0–0.33), mid (0.33–0.66), and outer (0.66–1.0)—revealed subtle regional differences. In the male kidney, average glomerular volumes across the three zones were 3.00 ×10^6^ ± 1.07 ×10^6^ *µm*^3^, 3.16 ×10^6^ ± 1.10 ×10^6^ *µm*^3^, and 3.27 ×10^6^ ± 1.06 ×10^6^ *µm*^3^, respectively. The female kidney exhibited slightly larger glomeruli across all regions, with values of 4.30 ×10^6^ ± 2.05 ×10^6^ *µm*^3^, 4.64 ×10^6^ ± 2.22 ×10^6^ *µm*^3^, and 4.55 ×10^6^ ± 2.16 ×10^6^ *µm*^3^, respectively. These regional patterns may hint at underlying physiological or pathological remodelling, particularly in the hypertensive donor Hoy et al. [2008].

## Discussion

### Model performance

The proposed correlative glomeruli segmentation pipeline evaluated four different deep neural networks on high-resolution, manually annotated data. nnUNet performed best and was therefore selected for correlative segmentation on lower-resolution data. This allows segmentations to be propagated from HiP-CT high-resolution VOIs to low-resolution complete organ scans.

HiP-CT is an inherently isotropic 3D modality, creating individual images that range in size from gigabytes to terabytes. As a result, training deep learning models on this data demands significant computational resources. The challenge is balancing the richness of contextual information in the inputs with the constraints of computational memory. Therefore, the training patches dimension was set to 128^3^ (filling 2GB of memory), which is large enough to contain complete glomeruli in high-resolution data while fitting into available memory for a batch size of two. The model performs best on the intermediate-resolution data, achieving a mean validation Dice score of 0.949, as more complete glomeruli with relatively clear boundaries are involved and more contextual information surrounding the glomeruli was provided in this resolution, compared to the other two resolutions, which have mean validation Dice scores of 0.860 and 0.796.

Contrast enhancement as a pre-processing step for HiP-CT data plays a key role in training a successful segmentation model. As shown by a recent Kaggle competition to segment kidney vasculature on HiP-CT data Jain et al. [2024], the ranges of voxel intensities are narrow and change between samples and resolutions due to factors including the X-ray scan setup, anatomical differences, and sample preparation, which are different for each sample and resolution. This results in the shift and bias of the model weights if the training dataset involves several samples. Therefore, we applied CLAHE, which can enhance local contrast while maintaining overall brightness, making small structures, such as glomeruli, more distinguishable (see Supporting Information S1 for more details). However, we used default hyperparameters when applying CLAHE to different samples. Although this helps generate a similar and wide range of voxel intensity for each sample, future works could explore the statistical impact of varying these parameters or investigate alternative contrast enhancement techniques applied to HiP-CT data. This will be particularly important for extending the model to larger cohorts of HiP-CT kidney data collected over recent years, during which the HiP-CT imaging methodology has evolved and data quality has significantly improved.

The proposed multiscale segmentation pipeline, whether applied to glomeruli as in this study or applied to other organ structures as a more general concept, can lead to cumulative errors introduced during the generation of pseudo-labels from higher-resolution predictions. The errors arise from several factors, such as segmentation inaccuracies in the model at previous scales, misalignments during multi-resolution registration, and the loss of structural details due to decreased resolution from the VOI scans up to the complete organ scans. As the pipeline progresses across scales, these errors can accumulate and become more noticeable, ultimately reducing the reliability of the segmentations at the lowest resolution. To address this, we developed an automated post-processing technique at each scale to eliminate false positives based on multiple criteria and validated against (pseudo-)ground truths.

In addition to post-processing, another key strategy for reducing error accumulation in the multiscale segmentation pipeline is improving imaging quality. All data used in this study were acquired on the beamline BM05 in the period from 2020 - 2022, before the start of the new BM18 beamline. The increased capabilities of beamline BM18, notably the increase in propagation distance for the lower resolution scans and the larger beam size, mean that the speed and contrast sensitivity of scans can be higher than those used to develop this model Vijayakumar et al. [2024]. For example, a whole kidney can be overviewed at 10 *µm*/voxel on BM18, or a 20 *µm*/voxel scan can be performed with 20 meters of propagation (as opposed to 3.5 meters), significantly enhancing the contrast sensitivity and thus clarity of the images. Using these scanning conditions on beamline BM18 enables higher throughput experiments, allowing multiple kidneys to be imaged in a single acquisition under more consistent image conditions. This reduces the sample-to-sample variability, simplifies the pre-processing requirements and improves the robustness of the segmentation model. An intact organ in a higher voxel size and better resolution helps reduce the number of resolution transitions within the multiscale segmentation pipeline. Indeed, we have already collected such imaging data and are actively applying the pipeline developed in this study to a larger number of studies to investigate general trends in glomerular morphology and spatial distribution.

Whilst higher imaging quality and consistency are always fundamental to the biological conclusions and applications for this multiscale segmentation, the challenge of feature degradation is also inherent to this approach. As resolution decreases across the hierarchical pipeline, image feature degradation limits the visibility of the fine features (as illustrated in Figure 1). In our case, this degradation affects glomeruli, but as the quality of images increases, the segmented target at the highest resolution might change to smaller structures, e.g. single cells. Despite better imaging quality, propagating segmentations of such fine structures across scales will face the same challenge. Therefore, segmentation accuracy can be compromised due to pixelation and the limitation of contrast sensitivity with lower-resolution data.

To mitigate these challenges, advanced feature enhancement such as a super-resolved tomographic reconstruction technique Krüppel et al. [2024] has been proposed. However, it is important to acknowledge that this multiscale segmentation approach cannot be infinitely propagated to lower-resolution data simply through image enhancement. At some point, the lower-resolution data lacks sufficient statistical information to support segmentations of specific features. This limitation is closely tied to the constraints of generative models and the challenge of hallucination Tivnan et al. [2024]. Fundamentally, the model presented in this study serves not only as a practical multiscale segmentation tool but also as a framework to evaluate when the feature can be confidently inferred from lower-resolution data through image enhancement and when higher-resolution imaging becomes necessary.

### Glomerular morphological analysis

The human kidney is traditionally thought to contain approximately one million glomeruli Kanzaki et al. [2015], though more recent studies have revealed substantial variability, with up to a 13-fold difference reported among healthy individuals Hoy et al. [2010]. Autopsy studies indicate that nephron number tends to be lower in hypertensive individuals compared to age-matched normotensive controls Keller et al. [2003]. Age-related nephron loss is also well-documented, with an estimated decline of ca. 3,676 glomeruli per kidney per year after age 18 Charlton et al. [2021]. In hypertensive patients, this loss may accelerate to ca. 200,000 glomeruli per decade after age 50, emphasising the combined impact of ageing and hypertension on nephron number Charlton et al. [2021].

The average glomerular volumes observed in our healthy male kidney align well with values reported by Denic et al. Denic et al. [2023] (2.9 × 10^6^ ± 1.1 × 10^6^ *µm*^3^, 3.2 × 10^6^ ± 1.2 × 10^6^ *µm*^3^, and 2.5 ×10^6^ ±1.1 × 10^6^ *µm*^3^ in juxtamedullary, mid-cortical, and superficial regions, respectively). Hughson et al. Hughson et al. [2014] reported that hypertension results in a 20–30% increase in glomerular volume, and our findings are consistent with this trend. In the hypertensive female donor, glomerular size was elevated across all cortical regions, which may suggest that hypertension induces global glomerular hypertrophy rather than region-specific effects.

Additionally, in our data, glomerular size showed a mild increase from the inner (near-medullary) to the outer cortical regions, especially in the healthy kidney. This spatial trend is consistent with previous observations by Terry et al. Samuel et al. [2005], who reported region-dependent variation in glomerular volume within the cortex. Regarding glomerular density, Kanzaki et al. Kanzaki et al. [2013] showed that it is highest near the superficial cortex and decreases with increasing distance from the renal capsule. Our model successfully captures this pattern by predicting a local peak near the cortex (Fig. 7 (C.1)). However, we also observed an additional peak in glomerular density near the medullary boundary. This secondary peak likely arises from the way relative distance is defined in our model, as glomeruli located between adjacent medullas have a large distance to the renal capsule and therefore appear artificially close to the medulla.

## Conclusion

Studies on hierarchical biomedical systems are critical to discovering fundamental physiological functions across multiple scales. This requires advanced imaging techniques to produce images across complete macro-scale organs and micro-scale near-cellular structures, and effective multiscale image processing pipelines to enable integrated analysis across these resolutions. In this work, we used HiP-CT, which can image intact human organs at ca. 20 *µm*/voxel down to high-resolution VOIs at ca. 1 *µm*/voxel. It inherently provides multiscale biomedical image datasets that capture the tiny functional structures at different resolutions. Using glomeruli segmentation in human kidney as a case study, we developed a hierarchical segmentation pipeline based on deep neural networks. Our results demonstrated that the segmentations can be propagated from high-resolution VOIs to entire low-resolution kidneys, enabling anatomical morphology analyses of those tiny functional units like glomeruli. While this work focused on human kidneys, the principles and architecture are generalisable to other organs that also exhibit similar multiscale structures involving small functional units across entire organs.

## Acknowledgments

This project has been made possible in part by grant number 2022-316777 from the Chan Zuckerberg Initiative DAF, an advised fund of Silicon Valley Community Foundation. The authors would also like to acknowledge ESRF beamtimes md1252, md1290, and md1389 as sources of the data, PDL is supported by Royal Academy of Engineering (CiET1819/10), we would also like to acknowledge EPSRC grant JADE-2 [EP/T022205/1].

