## supporting_information for "Multiscale Segmentation using Hierarchical Phase-contrast Tomography and Deep Learning"

### Supporting Information for: Multiscale Segmentation using Phase-contrast Tomography and Deep Learning

#### Contents

**1 HiP-CT tomographic information** 2

**2 HiP-CT training data** 3

**3 Data pre-processing** 4

**4 HiP-CT multiscale registration** 5

**5 Prediction post-processing** 6

**6 Training on complete organ scans** 8

**7 Results of LADAF-2021-17 right kidney** 10

### 1 HiP-CT tomographic information

Table 1 summarises the key scanning parameters for each sample and voxel size used in this study. The reconstructed datasets are available for download via the DOI links provided in the Dataset section of the main manuscript. Additionally, the portal **human-organ-atlas.esrf.eu** offers in-browser visualisation using Neuroglancer and access to supplementary data, including metadata, donor medical information, and image reconstruction parameters.

**Table 1.** HiP-CT scan parameters

| Organs | Voxel size ($\mu m$) | Data label | Acquisition mode | Projection number | Projection distance ($m$) | Attenuators | Average energy ($keV$) | Surface dose rate ($Gy/s$) | Number of scans | Per scan time ($min$) |
|---|---|---|---|---|---|---|---|---|---|---|
| S-20-28 | 2.5 | VOI-01,<br>VOI-02,<br>VOI-03,<br>VOI-04,<br>VOI-05,<br>VOI-06,<br>VOI-07, | Half | 6000 | 1.44 | Al 0.51mm,<br>Mo 0.24mm,<br>SiO2 16mm<br>rods 4x4mm | ~81 | U.N | 23,<br>17,<br>8,<br>5,<br>5,<br>9,<br>5 | U.N |
| LADAF-2020-17<br>Left Kidney | 2.6 | VOI-01.1,<br>VOI-02.1 | Half | 6000 | 1.44 | Mo 0.23mm,<br>SiO2 40mm<br>rods 10x4mm | ~83 | 35 | 15,<br>17 | 5.3 |
| LADAF-2020-17<br>Right Kidney | 2.6 | VOI-01.1,<br>VOI-02.1,<br>VOI-03.1 | Half | 6000 | 1.44 | Mo 0.23mm,<br>SiO2 40mm<br>rods 10x4mm | ~83 | 35 | 49,<br>8,<br>24 | 5.3 |
| | 6.5 | VOI-01,<br>VOI-02,<br>VOI-03, | Half | 6000 | 3.5 | Mo 0.23mm,<br>SiO2 40mm<br>rods 10x4mm | ~83 | 35 | 44,<br>13,<br>35 | U.N |
| | 25.0 | Complete organ | Half | 6000 | 3.5 | Mo 0.1mm,<br>SiO2 32mm<br>rods 8x4mm | ~81 | U.N | U.N | 4.3 |
| LADAF-2020-27<br>Left Kidney | 1.29 | Central column | Half | 6000 | 0.5 | Al 2mm,<br>Mo 0.1mm,<br>SiO2 bars<br>3*5mm | ~74 | 161 | 6 | 12 |
| | 6.05 | Central column | Half | 6000 | 3.475 | Al 2mm,<br>Mo 0.1mm,<br>SiO2 60mm<br>rods 12x5mm | ~89 | 10.5 | 13 | 5 |
| | 25.08 | Complete organ | Half | 6000 | 3.475 | Al 2mm,<br>Mo 0.1mm,<br>SiO2 60mm<br>rods 12x5mm | ~93 | 10.5 | 50 | 2.5 |

#### 2 HiP-CT training data

Fig. 1 illustrates the extracted 2D slices from the training data of LADAF-2020-27 left kidney. The 2D visualisations here are displayed at spatial dimensions of 512×512 for high resolution, 284×284 for intermediate resolution, and 657×657 for low resolution, respectively, preprocessed by 3D CLAHE before cropping as training patches.

During scanning, high-resolution volumes were normally selected from cortical regions, which are densely populated with glomeruli. These regions are within the intermediate-resolution column. To ensure effective threshold selection during the post-processing stage, particularly for false positive removal, it was necessary to introduce empty cubes (i.e., volumes without glomeruli) at both high and intermediate resolutions for effective validation.

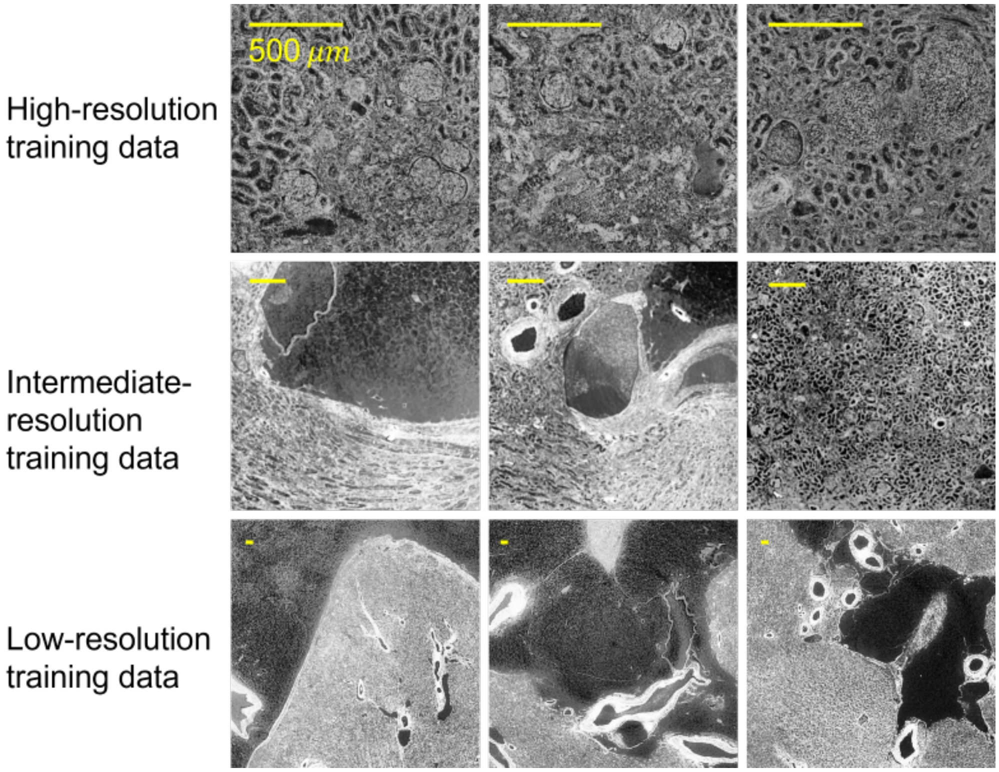


**Fig 1.** Training data visualised in 2D extracted from LADAF-2020-27 left kidney data imaged by HiP-CT. The high-resolution data (2.58 $\mu m$/voxel) and intermediate-resolution data (12.1 $\mu m$/voxel) do not involve much fat area, so they require additional empty cubes to evaluate the post-processing effects. However, the low-resolution data (25.08 $\mu m$/voxel) has a larger field of view.

### 3 Data pre-processing

Data pre-processing is a crucial step for preparing HiP-CT datasets before training neural networks. As illustrated in Fig. 2, the raw HiP-CT data are 16-bit with a narrow and sample-dependent intensity range. Additionally, training with 3D 16-bit volumes, even after cropping into smaller 3D patches, presents computational challenges. To address these issues, we applied 3D Contrast-Limited Adaptive Histogram Equalisation (CLAHE) to the 16-bit data volume first to enhance contrast and normalise intensity distribution. Then, the processed data were converted to 8-bit format, reducing memory requirements and improving training efficiency.

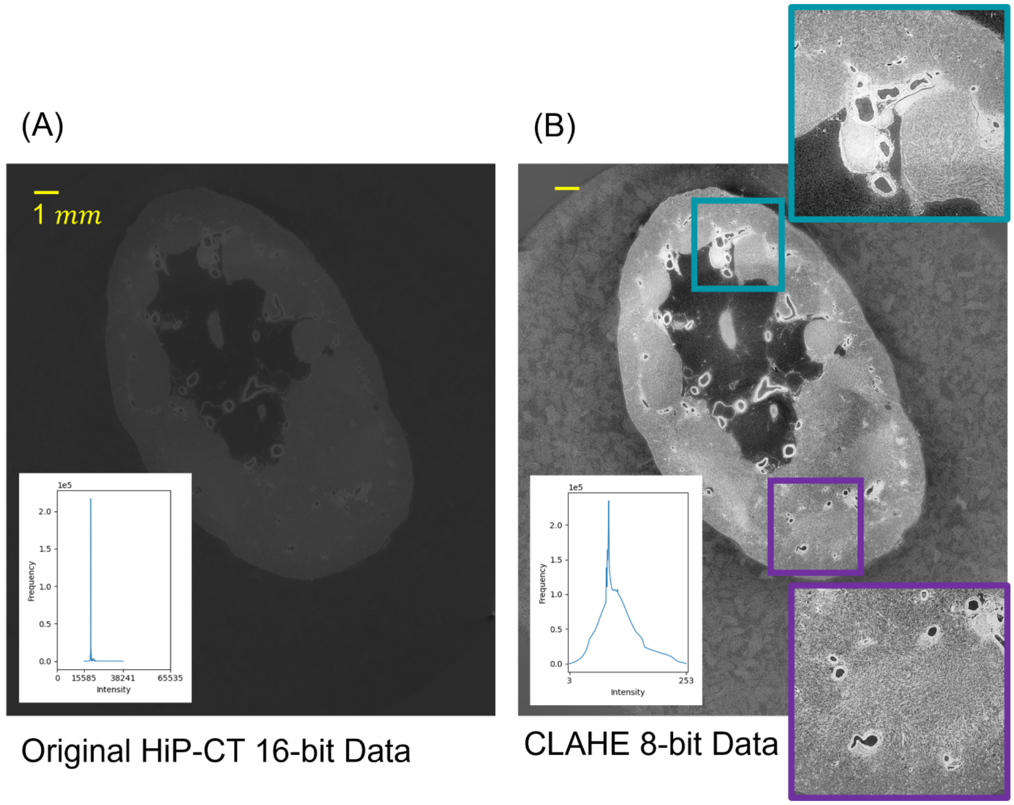


**Fig 2.** HiP-CT pre-processed by CLAHE and 8-bit conversion, compared to the original 16-bit HiP-CT data. The zoomed regions (blue and purple squares) show that the textures and features were highlighted after applying CLAHE.

### 4 HiP-CT multiscale registration

Multiscale registration is a key component of the proposed segmentation pipeline, enabling the generation of pseudo-labels and training datasets at lower resolutions. As shown in Fig. 3(A), a representative 2D slice from the complete kidney scan of LADAF-2020-27 at 25.08$\mu m$/voxel resolution is displayed. Fig. 3(B) illustrates the result of aligning this low-resolution slice as a fixed image with the corresponding higher-resolution counterpart as a moving image at 12.1 $\mu m$/voxel, demonstrating the effectiveness of the registration across scales.

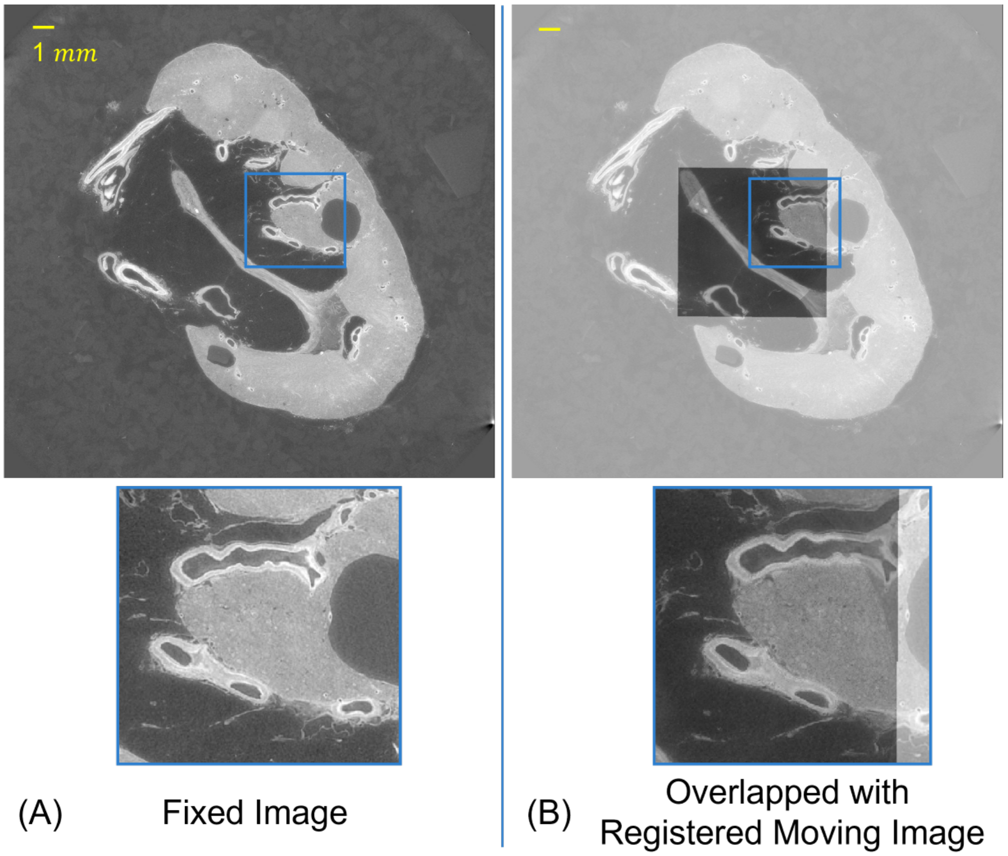


**Fig 3.** Registration between two resolutions of HiP-CT data. Samples were from LADAF-2020-27 left kidney, complete organs at 25.08 $\mu m$/voxel and intermediate-resolution column at 12.1 $\mu m$/voxel (binned by 2 from the original column at 6.05 $\mu m$/voxel)

### 5 Prediction post-processing

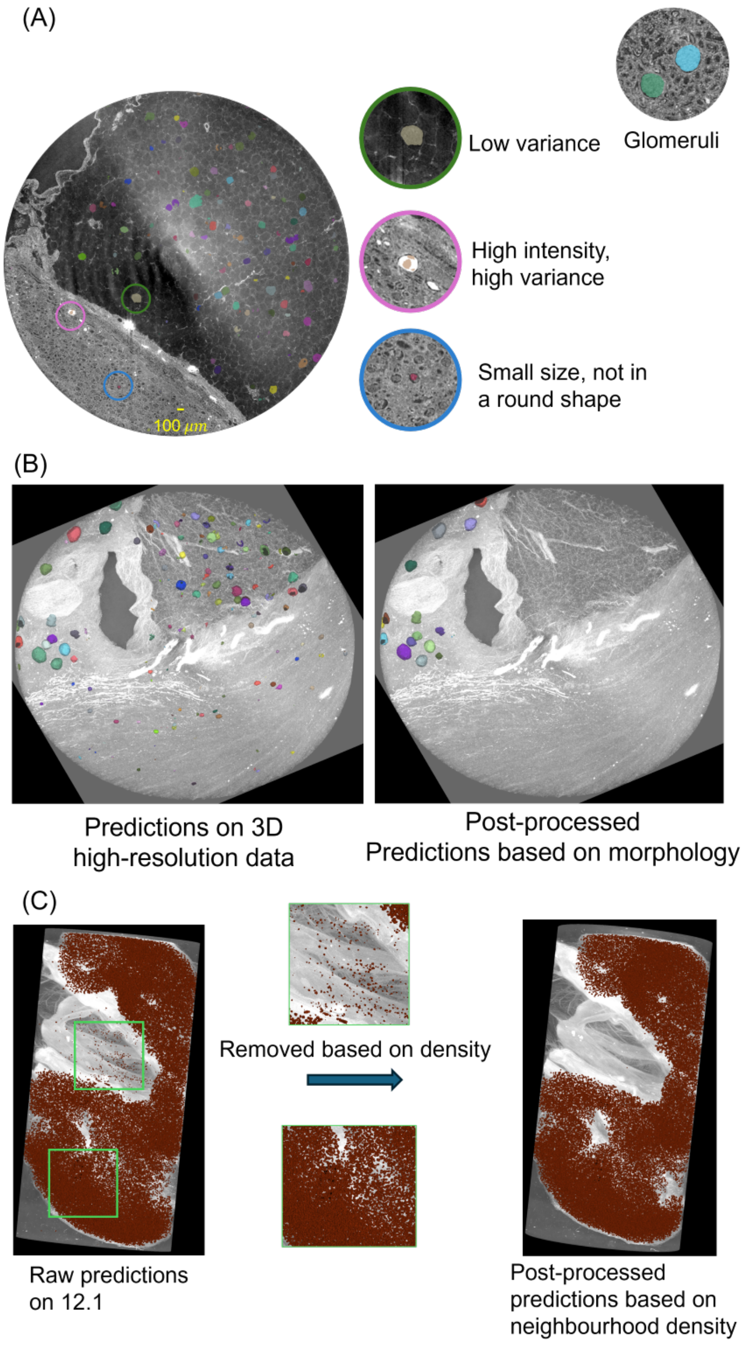


**Fig 4.** Effects of post-processing on segmentation predictions. (A) Examples of false positives in high-resolution data (LADAF-2020-27 Left Kidney, 2.58 $\mu m$/voxel) that can be eliminated using intensity-based and morphological criteria. (B) Segmentation result after applying post-processing to high-resolution data. (C) False positives in intermediate-resolution data (12.1 $\mu m$/voxel) that require an additional density-based filtering step for effective removal.

The effect of post-processing techniques is shown in Fig. 4. Panels (A) and (B) show results after post-processing on high-resolution data at 2.58 $\mu m$/voxel. At this resolution, false positives, classified into three categories as described in the main manuscript, can be effectively removed using the thresholding parameters based on intensity and morphological properties such as intensity variances and glomeruli size. However, for lower-resolution data, as shown in Fig. 4 (C), these parameters alone are insufficient. Therefore, an additional post-processing step based on prediction density is introduced to further reduce false positives.

Fig. 5 presents the average Instance Dice scores and average Dice scores for all training cubes, including one additional empty cube per resolution. To determine optimal post-processing parameters, 20 sets of thresholds were generated for each resolution using Latin Hypercube Sampling (LHS). Among the generated threshold sets with the same performance, the selection criterion is to preserve as many true positives as possible while eliminating false positives. For high-resolution data at 2.58 $\mu m$/voxel, the selected configuration (search No. 12) applied a broader intensity variance range and a lower roundness threshold and effectively preserved predicted glomeruli. In contrast, for the intermediate-resolution data at 12.1 $\mu m$/voxel, where only the lower bound of intensity variance was used to remove the fat area, the set of thresholds (search No.5) featuring the lowest variance and roundness was selected.

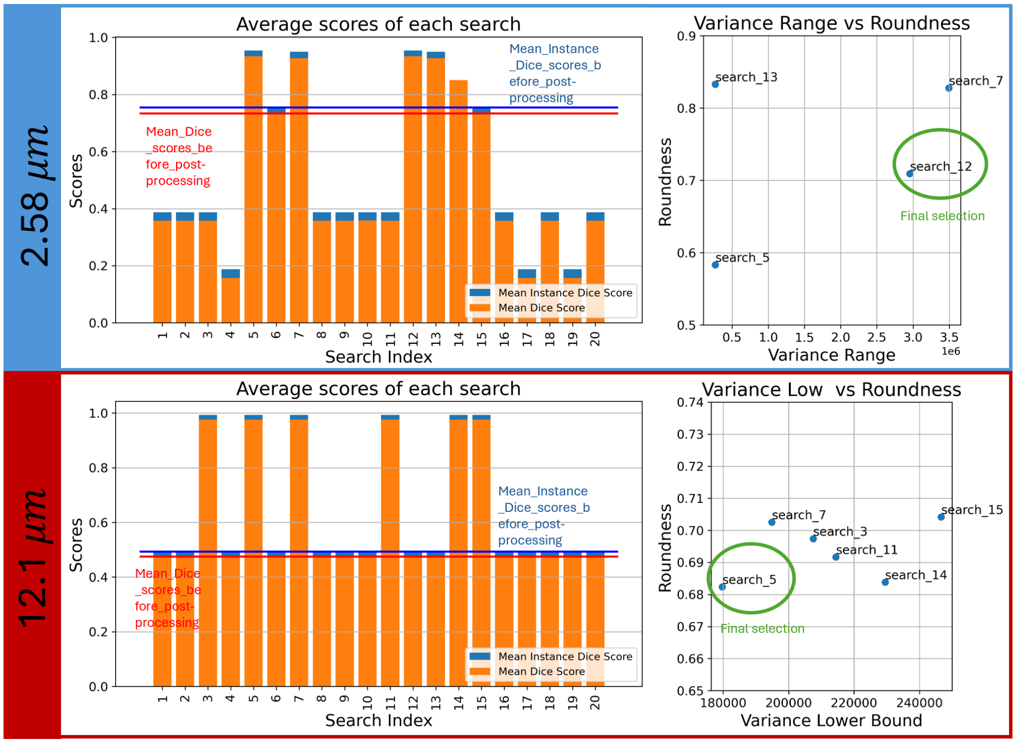


**Fig 5.** Selection of post-processing thresholds using Latin Hypercube Sampling (LHS). The bar chart displays the Dice scores and Instance Dice scores for each threshold set sampled via. LHS, validated by all the training cubes (involving an additional empty cube). The horizontal lines indicate the mean Dice score (in red) and mean Instance Dice score (in blue) across all the validation cubes. The scatter plot illustrates how the optimal parameter set was selected among those with the same performance, prioritising thresholds that preserve more true positives.

### 6 Training on complete organ scans

As discussed in the main manuscript, training on complete organ scans at low resolution (25.08 $\mu m$/voxel) is challenging due to the image degradation. Fig. 6 presents the original low-resolution HiP-CT 2D slice alongside the corresponding CLAHE enhanced slice and annotation from the training data of LADAF-2020-27 left kidney. Although CLAHE improves overall contrast and enhances visibility of structural boundaries, glomerular features remain substantially degraded at this resolution, making it challenging for the model to distinguish them reliably. Therefore, to improve the segmentation performance and the Dice scores, we investigated several training strategies as shown in Fig. 7.

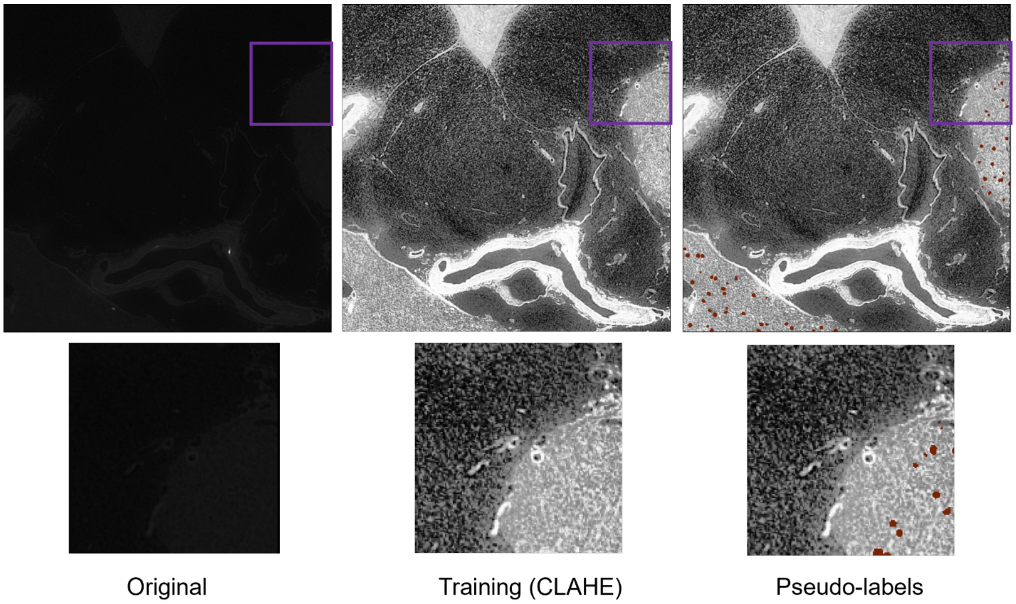


**Fig 6.** The 2D slices from complete organ scan of LADAF-2020-27 left kidney at 25.08 $\mu m$/voxel. The original image, the training image after CLAHE applied and the pseudo-labels from the predictions of the previous resolution in the multi-scale segmentation pipeline are shown.

We first compared fine-tuning versus training from scratch, as in panel (A) of Fig. 7. The results show that fine-tuning yields a higher Dice score after training 1000 epochs, indicating better performance. Given the low-resolution nature of the data, where pseudo-labelled glomeruli normally occupy a small fraction of the volume, we also sought to reduce background dominance. To do so, we excluded 3D patches in which glomerular labels comprised less than 1% of the volume. Panel (B) shows a comparison between training on the full dataset versus the filtered datasets without low-label patches over 1000 epochs. We found that removing these nearly empty cubes significantly improved model performance. Given the similar performances between the trainings on the datasets of removing all the cubes with labels volume smaller than 1%, and keeping 0.7% of those cubes, we used the latter for training on the low-resolution whole kidney data.

Despite these improvements, Dice scores seemed to continue to rise beyond 1000 training epochs, indicating under-convergence. Therefore, we explored extended training epochs with different learning rate scheduling, as shown in panel (C). Two learning rate decay strategies were tested: polynomial decay (Eq. 1) and exponential decay (Eq. 2):

$$LR_{poly} = \begin{cases} 0.01 \times (1 - \frac{x}{1000})^{0.9}, & \text{if } 0 \leq x < 1000, \\ 0.002 \times (1 - \frac{x}{1500})^{0.9}, & \text{if } 1000 \leq x < 1500, \end{cases} \tag{1}$$

$$LR_{exp} = 0.01 \times 0.994^{x}, \tag{2}$$

where $LR_{poly}$ and $LR_{exp}$ denotes the learning rate at epoch $x$ for polynomial and exponential decay, respectively. Considering that the Dice score increased very slowly around 1000 epochs in previous experiments, both polynomial and exponential learning rate schedules were designed to be small when continuing training from 1000 epochs to avoid the overfitting problem. While exponential decay accelerated convergence, polynomial decay consistently led to better segmentation performance. Additionally, extending training to 1500 epochs provided only a slight improvement, suggesting that further training beyond this point would likely involve a tradeoff between performance gain and computation cost. Therefore, the final model for low-resolution data was trained using polynomial learning rate decay over 1500 epochs.

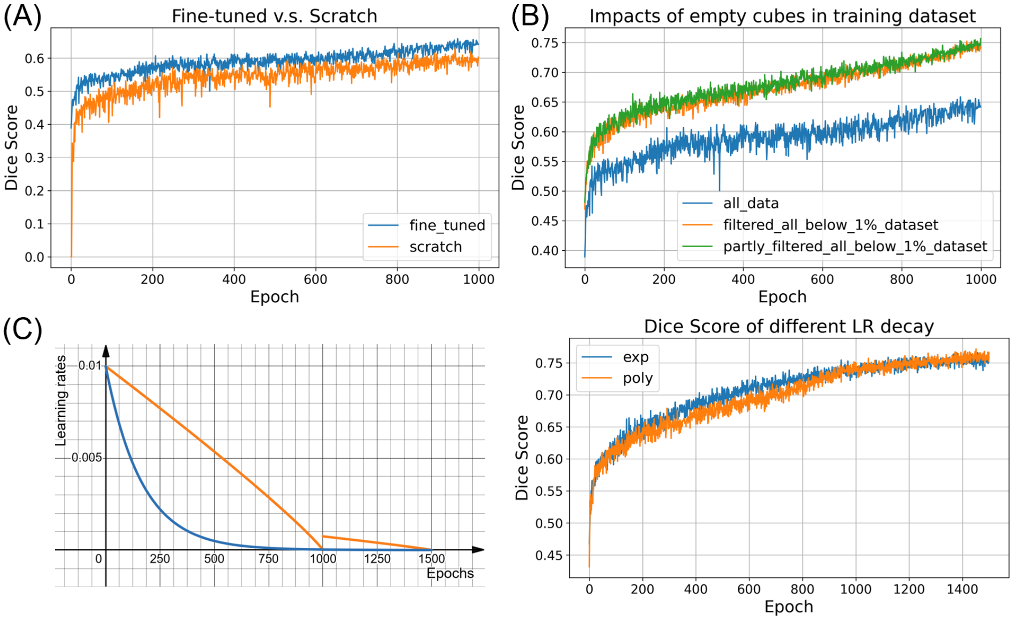


**Fig 7.** Evaluation of different training strategies for low-resolution complete organ scans at 25.08 $\mu m$/voxel.(A) Comparison between a fine-tuned model initialised from intermediate-resolution training and a model trained from scratch using nnUNet defaults. (B) Performance comparison between training on all registered cubes (blue line), on a subset with all the cubes with label volume ¡1% filtered (orange line), and a subset keeping 0.7% of the cubes with label volume ¡1% (green line). (C) Impact of learning rate schedules: polynomial decay vs. exponential decay.

### 7 Results of LADAF-2021-17 right kidney

This section presents the multiscale segmentation results for the LADAF-2021-17 right kidney, which was used for morphological analysis. For this kidney, the intermediate-resolution volumes were at 13 $\mu m$/voxel and the low-resolution complete kidney was acquired at 25 $\mu m$/voxel. We applied the high-resolution model trained on manually annotated data to the VOI-02 and VOI-03. These volumes were subsequently registered to the low-resolution complete kidney volume to generate the training data for fine-tuning the model.

As described in the main manuscript, 5-fold cross-validation of nnUNet on the LADAF-2020-27 left kidney did not show significant variation across folds. Therefore, for LADADF-2021-17 right kidney, we only implemented fold 0 training and used the model for glomeruli segmentation to streamline the workflow. Table 2 shows the number of training and validation patches used for fine-tuning at each resolution, with a 9:1 train/validation split. This table also reports the best Dice score achieved during the fine-tuning processes. Fig. 8 further shows the detailed training curves.

**Table 2.** LADAF-2021-17 Right Kidney training dataset sizes and best Dice scores during training for each resolution.

| | Dataset size (patches) | | Best Dice scores | |
|---|---|---|---|---|
| | Training | Validation | Training | Validation |
| 13 $\mu m$/voxel | 1498 | 375 | 0.935 | 0.915 |
| 25 $\mu m$/voxel | 1556 | 390 | 0.803 | 0.805 |

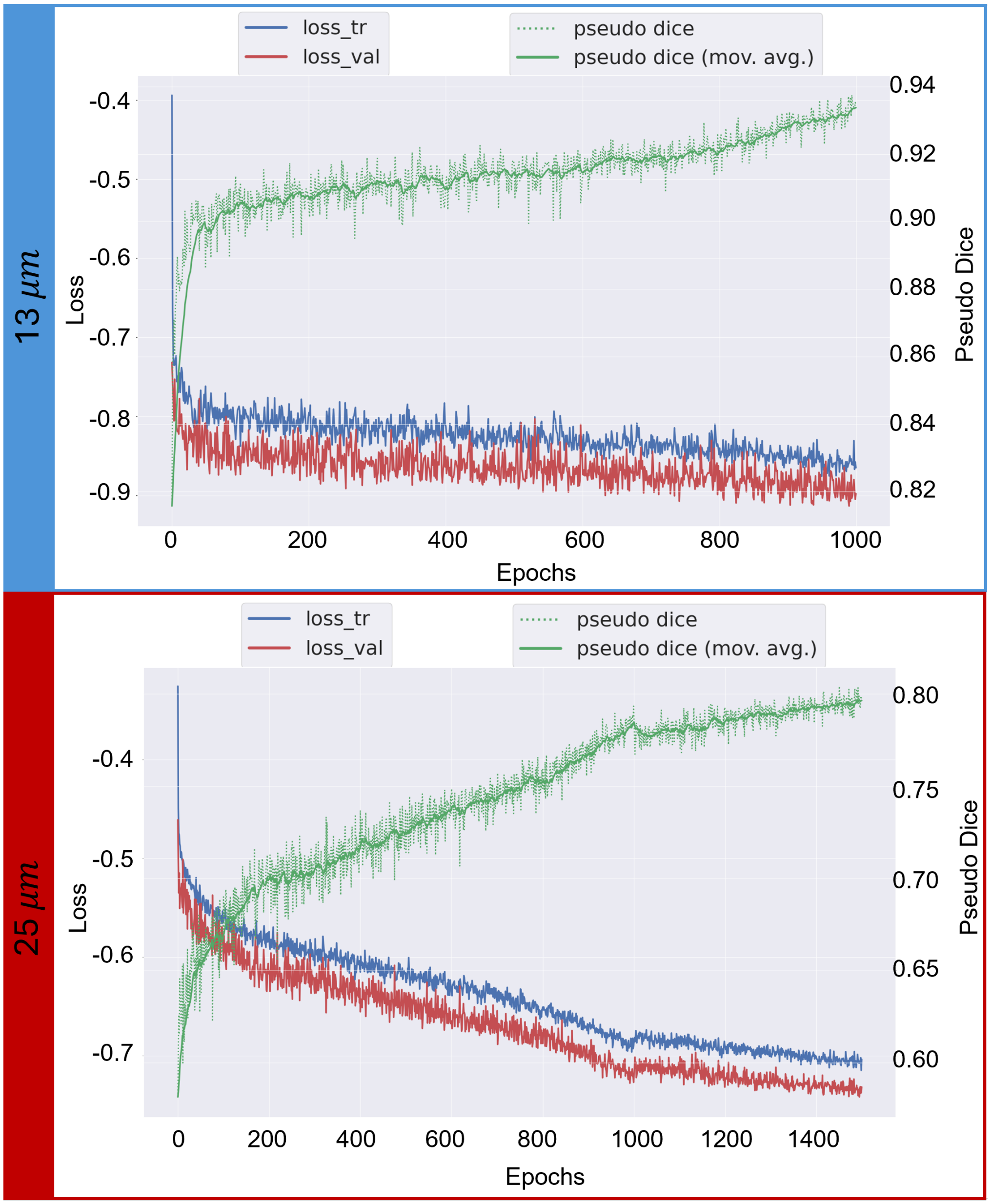


**Fig 8.** Curves of training losses and pseudo Dice scores at each resolution fine-tuning for the LADAF-2021-17 right kidney sample.


#### Acknowledgments

This project has been made possible in part by grant number 2022-316777 from the Chan Zuckerberg Initiative DAF, an advised fund of Silicon Valley Community Foundation. The authors would also like to acknowledge ESRF beamtimes md1252, md1290, and md1389 as sources of the data, PDL is supported by Royal Academy of Engineering (CiET1819/10), we would also like to acknowledge EPSRC grant JADE-2 [EP/T022205/1].
